# Selective infection and loss of PRDM1+ LN Tfh cells in uncontrolled HIV infection precludes formation of Tfh reservoirs under ART

**DOI:** 10.64898/2025.12.11.693731

**Authors:** Vincent H. Wu, Jayme M.L. Nordin, M. Betina Pampena, Suzanna Rachimi, William L. Burgess, Michael Clark, Jake A. Robinson, Perla M. del Rio Estrada, Mariane H. Schleimann, Fernanda Torres-Ruiz, Mauricio González-Navarro, Gonzalo Salgado, Yara Andrea Luna-Villalobos, Gustavo Reyes-Terán, Katharine J. Bar, Jacob D. Estes, Ole S. Søgaard, Neil Romberg, Santiago Ávila-Ríos, Michael R. Betts

**Author notes:** Correspondence to: Dr. Michael R. Betts, PhD Department of Microbiology University of Pennsylvania | Dr. Vincent H. Wu, PhD Vaccine and Immunotherapy Center The Wistar Institute. These authors contributed equally to this work (Vincent H. Wu, Jayme M.L. Nordin).

## Abstract

The multifaceted and long-lived nature of HIV-1 reservoirs has proven to be a formidable obstacle in the development of a therapeutic cure of HIV-1. One of the major dimensions of the HIV-1 reservoir is its prevalence and persistence in tissue environments, including in lymph nodes (LN). Within LN, T follicular helper (Tfh) cells are widely considered as the dominant sub-reservoir in viremic people with HIV (PWH). However, whether Tfh cells survive to establish a reservoir in PWH undergoing suppressive antiretroviral therapy (ART) has remained controversial. To address these issues, we deeply phenotyped over 500,000 cells and identified over 2,000 HIV-1 infected cells by employing single-cell multiomics on LN from PWH during viremia and suppressed on ART. While we detected HIV-1 infected Tfh cells, the majority of infected cells during viremia and ART had non-follicular phenotypes and were instead heterogeneously distributed between various CD4+ T-cell subsets. Within-subset comparisons of HIV+ and HIV- cells revealed heightened activation signatures and altered cell cycle states during viremia, but largely similar features during ART. Furthermore, the comparison between viremia and ART in PWH highlighted a noncanonical Tfh subset - defined by high locus accessibility and transcription of *PRDM1* - that is selectively depleted during viremia, recovers after ART, and is highly susceptible to infection in vitro. Our work suggests a revised direction for the HIV-1 cure field where a mandate of any comprehensive strategy will be to address the high proportional burden of non-follicular cells within the HIV reservoir.

## Introduction

HIV-1 cure strategies require a comprehensive understanding of the HIV reservoir for effective and selective targeting. Even after initiation of combinatorial antiretroviral therapy (ART) that leads to undetectable viral loads, this pool of infected cells can remain present in people with HIV (PWH) and reactivate towards viral rebound after ART interruption^1–4^. Tissue compartments, especially the gut and lymph nodes (LN)^5–7^, are sites where the HIV reservoir is seeded and is persistent. As such, a complete cure strategy must address the HIV reservoir, whether it be elimination of infected cells or durable control against rebound. To do so, a deep immunological and phenotypic understanding of the tissue reservoir is critical to provide clarity on the reservoir composition in order to design specific and comprehensive HIV cure strategies.

Studies from our group and others have demonstrated the extensive heterogeneity of the HIV reservoir within peripheral blood, lymph node, and gut mucosal CD4+ T cells of viremic PWH using DNA, RNA, and/or protein-based single cell methodologies^8–12^. These studies have broadly demonstrated T cell subset-specific characteristics of HIV-infected CD4+ T cells, but have not yet revealed any specific signatures of infected cells (as compared to uninfected cells of the same subset) beyond the presence of the proviral genome, viral transcripts, or viral proteins. With these previously published studies, it is clear that the environment and immune composition of cells (regardless of infection) are starkly different between tissue and peripheral blood. Furthermore, in LN, acute and chronic HIV infection are known to alter the tissue environment with the hallmarks of follicular hyperplasia and/or fragmentation and persistent generalized lymphadenopathy^13,14^. Given the complexity of LN and their importance as a hub for adaptive immunity, our understanding of the HIV reservoir within LN requires an unbiased assessment to untangle the cellular heterogeneity during untreated and treated HIV infection.

Early studies have established that HIV replication predominately occurred in lymph node B cell follicles during untreated viremia, with subsequent works defining the infected cells in lymphoid follicles as CD4+ T follicular helper (Tfh) cells^15–18^. Given the strong preference for HIV to infect and replicate in activated cells^19,20^, it is unsurprising that CD4+ Tfh cells would be highly involved in HIV pathogenesis, as Tfh cells are amongst the most activated CD4+ T cells in LN during primary and ongoing immune responses. We have also observed infected Tfh cells in LN during chronic HIV with our earlier study, though HIV DNA was detected in other CD4+ T cells as well^9^. However, the contribution of CD4+ Tfh cells to the lymph node HIV reservoir during ART is less clear, considering the decrease in immune activation^21,22^. Of note, Perreau and colleagues found that Tfh cells harbored similar levels of HIV DNA compared to other CD4+ T cell types in the lymph node during ART^16^, while Banga and colleagues showed an enrichment of HIV DNA in PD1+ CD4+ T cells compared to CD4+ PD-1- cells during ART^17^. However, the expression of PD-1 in lymph node CD4+ T cells is widespread and not exclusive to Tfh cells^9,23^. As such, there remains limited evidence for an HIV reservoir that is predominantly located in the B cell follicles during ART.

To address this critical gap in knowledge of the lymphoid reservoir in a comprehensive manner, we profiled memory CD4+ T cells from a large number of lymph node samples during viremia and during ART using scDOGMAseq^24,25^ for the simultaneous assessment of epigenetics, transcriptomics, and surface antigen levels. Using the presence of transposed HIV-1 DNA during the epigenetic profiling and/or HIV RNA during transcriptomic profiling, we found HIV-1 infected cells and determined their cellular phenotype. Along with orthogonal validation, we found that the predominant infected cell type during viremia or during ART does not have a classical Tfh phenotype. Though they were not the majority of infected cells, Tfh cells were still important as we found specific depletion of a particular Tfh subset during viremia and identified a strong indicator for HIV infection. Our efforts to delve into the lymphoid tissue reservoir has generated valuable insights to aid in design of comprehensive HIV cure strategies where 1) there is no evidence of a specific cellular signature outside of the integrated provirus itself that would allow a universal targeted approach for selective elimination of the HIV LN reservoir and 2) cure efforts should not solely focus on Tfh reservoirs, as lymph node follicles may not harbor the majority of infected cells within lymphoid tissues, particularly during long-term ART.

## Results

### Single-cell DOGMAseq atlas of the lymphoid tissue CD4+ T cells from viremic and treated PWH

To establish a comprehensive and trimodal atlas of HIV+ cells from lymphoid tissue, we performed scDOGMAseq (**Supplemental Figure 1A**) on sorted CD4+ T cells from inguinal or cervical LN of people without HIV (n = 4), viremic HIV (n = 12), and on ART (n = 10). We sorted live, CD3+ CD8- non-naive T cells **(Supplemental Figure 1B)** using a low stringency memory gate to improve cell numbers for single-cell processing while allowing recovery of transitional memory cells or those in process of differentiation. After alignment to a chimeric human/HIV-1 genome (HXB2 strain) and filtering for quality control, we obtained a final dataset of 559,760 cells across the 26 tissue donors (**Table 1 and Figures 1A-B**) with 18.3% of the dataset from HIV- donors, 29.1% of the dataset from viremic donors, and 52.6% of the dataset from donors on ART. The latter population was intentionally collected at a higher proportion in order to improve the odds of infected cell recovery based on the expected lower reservoir size during suppressive ART^26,27^.

**Figure 1:**
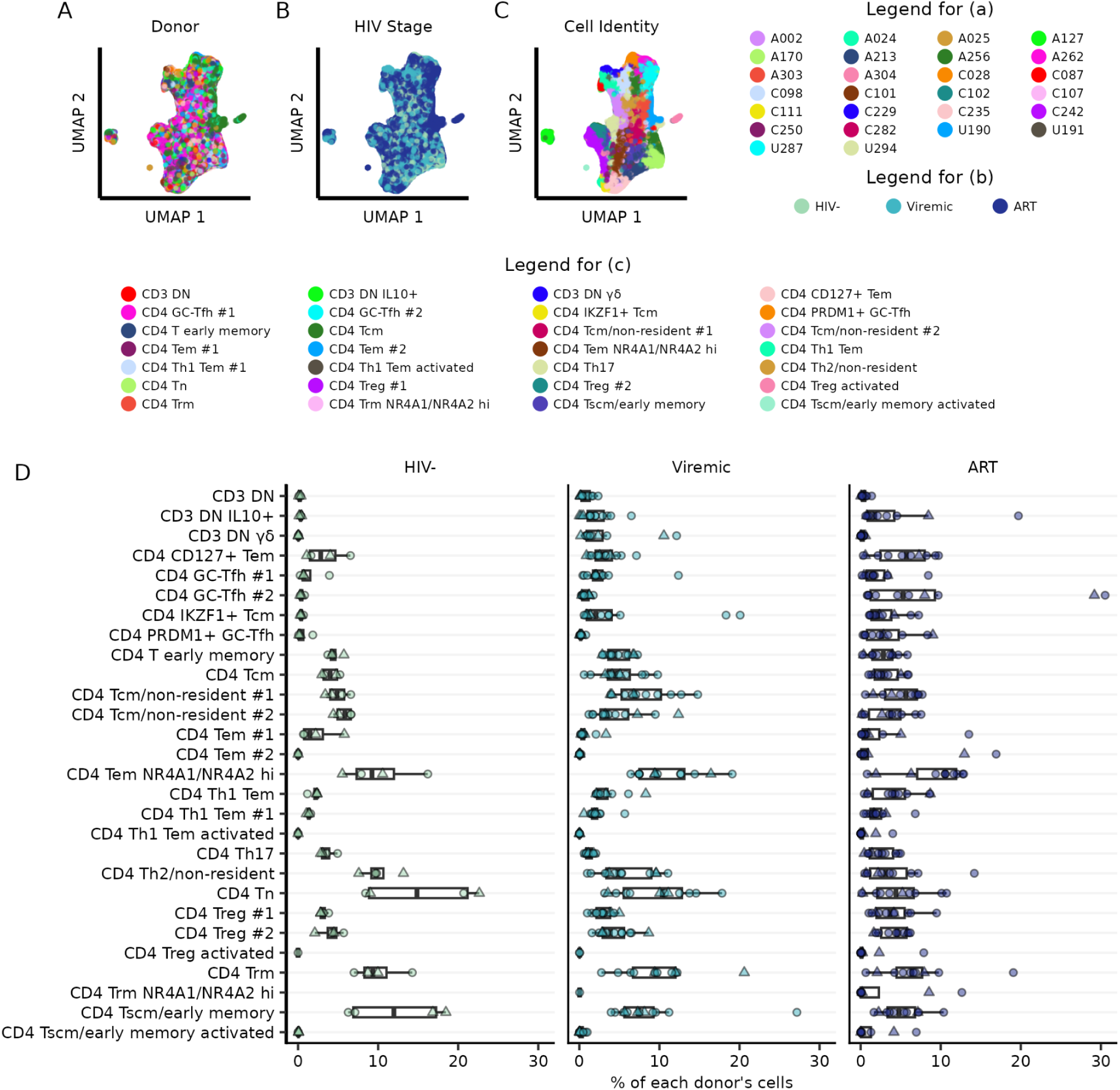
Single-cell DOGMAseq atlas of the lymphoid tissue from viremic and treated PWH. UMAP representations built on epigenetic and transcriptomic modalities of the scDOGMAseq dataset. Each point represents a cell and is colored either by (A) donor, (B) HIV infection stage, or (C) manual annotation. (D) Percentage of cells by donor (separated by HIV infection stage) that are found in the different, manually annotated cell clusters. Shape of dots indicate cervical (circle) or inguinal (triangle) LN.

Using unbiased clustering from an integrated space of epigenetic and transcriptomic dimensions, we identified and manually annotated 28 unique clusters based in part on a carefully curated reference list (**Supplemental Figure 2, Figure 1C, and Supplemental Table 1**). Where ambiguity between closely related clusters was apparent, we included immunologically related differentially expressed genes (DEGs) or open loci, such as “PRDM1” for the PRDM1+ GC-Tfh cluster, within the annotation. The annotated clusters consisted mostly of different memory CD4+ T cell subsets, but included some naive and innate-like lymphoid cells due to our more flexible gating strategy. While there was immense heterogeneity at inter- and intra-personal levels, several populations stood out with differential abundance (**Figure 1D**). These included two different subsets of CD4+ germinal center T follicular helper (GC-Tfh) cells that were enriched in donors on ART over viremic donors: a 13-fold difference (adj p = 0.067) for CD4 GC-Tfh #2 and a 15.7-fold difference (adj p = 0.003) for CD4 PRDM1+ GC-Tfh cells.

Furthermore, we observed a 2.84-fold depletion (adj p = 0.08) of Th17 cells in viremic donors over HIV- donors in alignment with previous studies^28–30^.Together, these data highlight the diversity of CD4+ T cell subsets in lymphoid tissue within and between individuals and their dynamic nature over the course of treated and untreated HIV infection.

### Dominant contribution of non-Tfh cell subsets to HIV reservoir during viremic and treated HIV-1 infection

To identify cells harboring viral RNA and/or DNA, we next employed previously established stringent detection thresholds for viral RNA (≥2 HIV related UMI or ≥4 reads of a single UMI/cell) and proviral DNA (≥1 UMI/cell)^9,10^. In viremic individuals, we found 1,795 HIV+ cells of which 1,183 (65.9%) had only HIV DNA, 454 (25.3%) had only HIV RNA, and 158 (8.8%) had both HIV DNA and RNA (**Figures 2A-B and Supplemental Figure 3A**). In individuals on ART, there were 400 HIV+ cells consisting of 306 HIV DNA+ (76.5%), 65 HIV RNA+ (16.3%), and 29 HIV DNA+ RNA+ (7.3%) cells (**Figures 2A-B and Supplemental Figure 3B**). Similar to a prior study on HIV+ cells from peripheral blood, the detection of HIV RNA+ only cells was not surprising due to the numerical odds of recovering high copies of HIV RNA compared to a single-copy ∼10kb viral DNA amongst the ∼3gb human genome^10^. Additionally, coverage across the reference viral genome indicated more enrichment of LTR reads during ART compared to during viremia (**Supplemental Figure 3C**), potentially indicative of defective and/or heterochromatic proviral integration. It is likely that the HIV RNA+ only cells also contain HIV DNA and as such, we binned the HIV RNA+ only cells with the dual detected cells (HIV DNA+ RNA+) in downstream analysis, which combined are labeled as HIV DNA+/- RNA+ for stringency.

**Figure 2:**
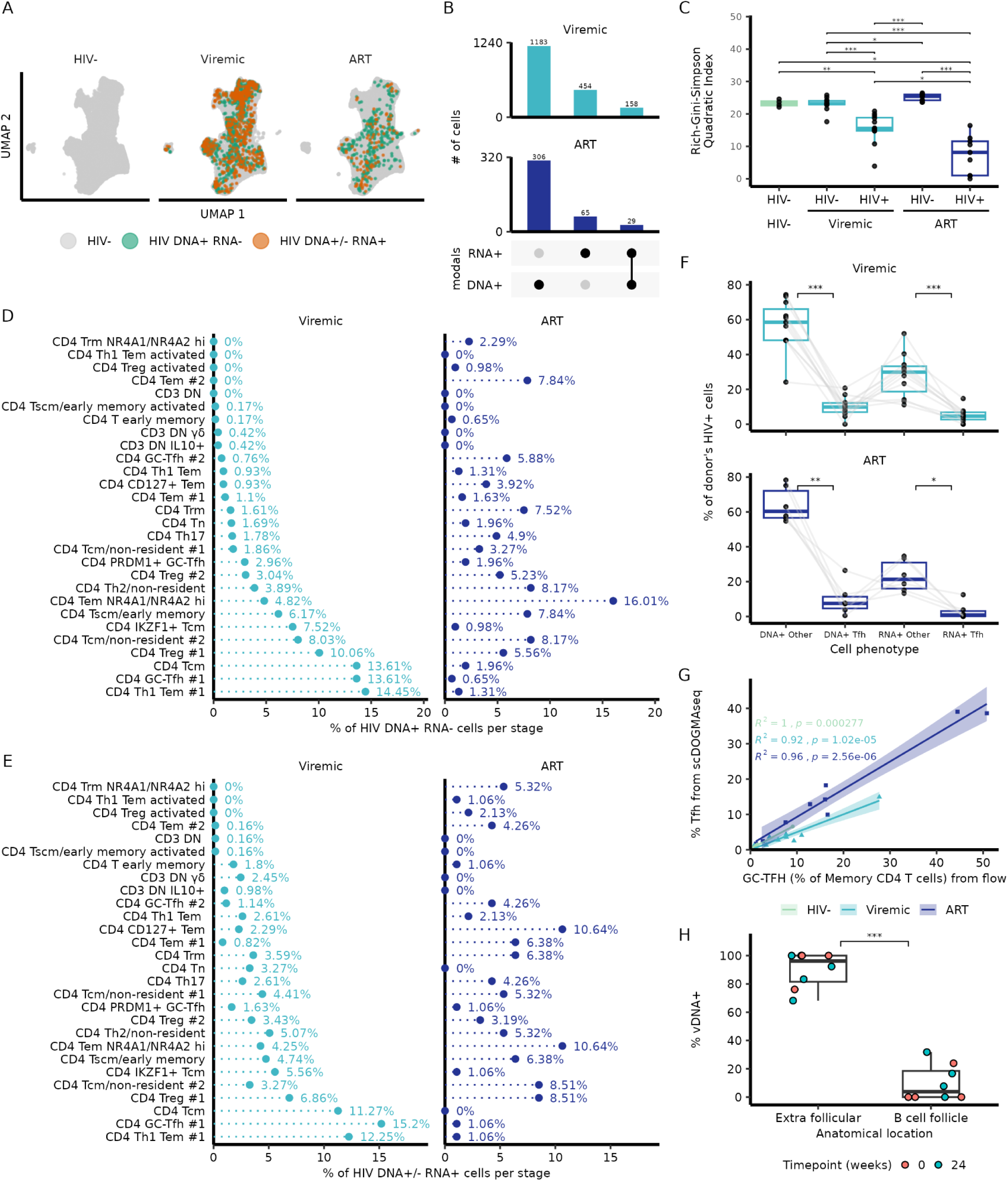
HIV+ cells are not predominantly in the Tfh compartment by scDOGMAseq. (A) UMAP representation in the same coordinate space as Figure 1, separated by infection stage. Cells are colored by detection of HIV DNA and/or RNA. (B) Quantification of the number of cells with HIV DNA and/or RNA detection across viremic individuals or ART-treated individuals. Cell counts are mutually exclusive (i.e. HIV DNA+ RNA+ cells are not counted extra in the single HIV DNA+ or HIV RNA+ category). (C) Weighted beta diversity is shown between HIV- and HIV+ cells separated by infection stage. P-values were calculated with the Wilcoxon rank sum test and corrected for multiple tests using the Holm method. (D) Distribution of all aggregated HIV DNA+ RNA- cells (separated by infection stage) across annotated clusters. Clusters are arranged in ascending order during viremia. (E) Distribution of all aggregated HIV DNA+/- RNA+ cells (separated by infection stage) across annotated clusters. Cluster order is the same as Figure 2D. (F) Percentage of donor’s HIV DNA+ RNA- (labeled as DNA+) or HIV DNA+/- RNA+ (labeled as RNA+) cells were binned by cell subset. The two bins were either Tfh (CD4 GC Tfh #1, CD4 GC Tfh #2, or CD4 PRDM1+ GC-Tfh) or “Other” (any other subset). Gray lines indicate the same donor. Percentages were compared using a paired Wilcoxon Rank Sum test with Bonferroni multiple test adjustment. (G) GC-Tfh (CD4 GC Tfh #1, CD4 GC Tfh #2, or CD4 PRDM1+ GC-Tfh clusters) proportions by donor were calculated from scDOGMAseq and compared with autologous GC-Tfh percentages (of memory CD4 T cells) measured from flow cytometry. Line shading indicates standard error with a linear model fit. R squared and p values were calculated with a two-sided Pearson correlation test. (H) Quantification of HIV vDNA+ cells that were found using histology and DNAscope from the LN acquired as part of the TEACH-B cohort. Points are colored by whether the sample was acquired before or after MGN1703 treatment. Comparisons were done using a Wilcoxon Rank Sum test with Bonferroni multiple test adjustment. * = adjusted p < 0.05; ** = adjusted p < 0.01; *** = adjusted p < 0.001

With robust detection of HIV+ cells in our dataset, we next examined the subset distribution of infected cells during viremia and ART. First, we assessed the diversity index by calculating the beta diversity via the Rich-Gini-Simpson Quadratic Index^31^. During viremia, HIV+ cells exhibited reduced diversity compared to HIV- cells (**Figure 2C**). The decrease in beta diversity also was observed during ART as well, suggesting that the HIV+ cell subset distribution is significantly different than HIV- cells.

We next asked what is the proportion of infected cells across viremic and PWH on ART. While the infected cells were heterogeneous and found in most cell clusters, we observed that the HIV DNA+ RNA- and HIV DNA+/- RNA+ cells were disproportionately distributed amongst cell clusters and differed by infection stage (**Figures 2D-E**). During viremia, the top two HIV DNA+ RNA- cell clusters included a GC-Tfh subset (CD4 GC-Tfh #1; 13.61% of all HIV DNA+ RNA-cells) and a Th-1 biased effector memory T cell (Tem) subset (CD4 Tem Th1 #1; 14.45% of all HIV DNA+ RNA- cells) (**Figure 2D)**. These clusters also dominated the distribution of HIV DNA+/- RNA+ cells from viremic individuals (15.2% and 12.25% of all HIV DNA+/- RNA+ cells, respectively) (**Figure 2E).** However, this distribution was dramatically shifted in ART-treated PWH with only 0.65% of HIV DNA+ RNA- cells being in the same Tfh subset and 1.31% of HIV DNA+ RNA- cells in the Tem Th1 subset. The largest cell subset that contributed to the pool of HIV DNA+ RNA- cells was instead recently activated Tem cells (CD4 Tem NR4A1/NR4A2 hi; 16.01% of HIV DNA+ RNA- cells from treated individuals) (**Figure 2D**). We observed a similar pattern for HIV DNA+/- RNA+ cells in treated PWH with recently activated Tem cells (CD4 Tem NR4A1/NR4A2 hi; 10.64% of all HIV DNA+/- RNA+ cells) and Tem cells expressing CD127 (CD4 CD127+ Tem; 10.64% of all HIV DNA+/- RNA+ cells) (**Figure 2E**).

Although the GC Tfh #1 cluster was one of the largest proportional contributors to HIV DNA+ or HIV RNA+ cells in viremic PWH, the vast majority (>70%) of infected cells were significantly skewed towards non-Tfh cell types in both viremic and treated PWH (threshold ≥10 HIV+ cells/cluster) (**Figure 2F**). To validate our detection of Tfh cells by scDOGMAseq, we independently assessed the proportion of CD4+ GC-Tfh (CXCR5+ PD1hi) cells on all available samples (n = 23). We found a significant and strong correlation (R^2^ > 0.9) between the detection of GC-Tfh cells by flow cytometry versus scDOGMAseq from people without HIV, viremic PWH, and ART-treated PWH (**Supplemental Figure 3D and Figure 2G**).

To further validate our findings in a different cohort with an orthogonal approach, we performed HIV DNAscope on 8 iliac LN sections from PWH on ART (one baseline and one post 24-weeks treatment, n=4 donors) in the TEACH-B cohort^6,32–35^. The TEACH-B clinical trial recruited PWH with at least 12 months of suppressive ART to study the impact of 24 weeks of TLR9 agonist (MGN1703) treatment. Similar to the previously reported levels of HIV-1 DNA remaining consistent before and after MGN1703 treatment^33^, the proportion of viral DNA+ cells from these tissue sections did not significantly change between the baseline measurement and 24 weeks of MGN1703 treatment (**Supplemental Figure 3E**). In strong congruence with our DOGMAseq data on ART-treated PWH, the percent of HIV DNA+ cells found within extra-follicular regions (median of 96.15%; range: 68.16-100%) was significantly higher than the percent found within B cell follicles (median of 3.85%; range: 0-31.82%) (**Figure 2H and Supplemental Figure 3F**) at baseline) and after 24 weeks of MGN1703. These data, from both DOGMAseq and DNAscope from an independent HIV cohort, indicate that while GC Tfh CD4+ T cells have a high proportional burden of infection in the absence of therapy, the majority of infected LN CD4+ T cells do not have a GC-Tfh signature and are not restricted to B cell follicles, especially during suppressive ART.

### Heightened T-cell activation signatures in HIV+ cells during viremia

Given that immune states are profoundly different during untreated viremic infection and during ART-treated viral suppression^36,37^, we sought to separate our analyses by infection stage. First, we specifically investigated if LN HIV+ cells differed from HIV- cells in viremic PWH. Using an unbiased approach to detect differential transcription factor motifs within accessible chromatin^38^ that were enriched in subset-specific HIV+ or HIV- cells compared to all other cells, we observed that HIV+ cells were overall more activated via the TCR receptor pathway compared to HIV-cells. This was seen with 1) increased enrichment of AP-1 family transcription factor motifs (Module 12); 2) NFATC3/NFACTC2 motifs (Module 10); and 3) NFKB1, NKFB2, and its related transcription factor motifs (Module 11) (**Figure 3A**). In concordance with responses towards viral infections^39^, there was increased enrichment in interferon-related transcription factors (Module 9) in HIV+ cells compared to HIV- cells. This effect, which was more pronounced in non-Tfh phenotypes (**Figure 3A**), is in agreement with previous studies that have shown antagonistic roles of STAT3 and type I interferon signaling in Tfh differentiation^40^. We also observed greater enrichment of POU family transcription factor motifs in HIV+ cells (Module 9; **Figure 3A**) that are known to be important for HIV replication and infection^41^.

**Figure 3:**
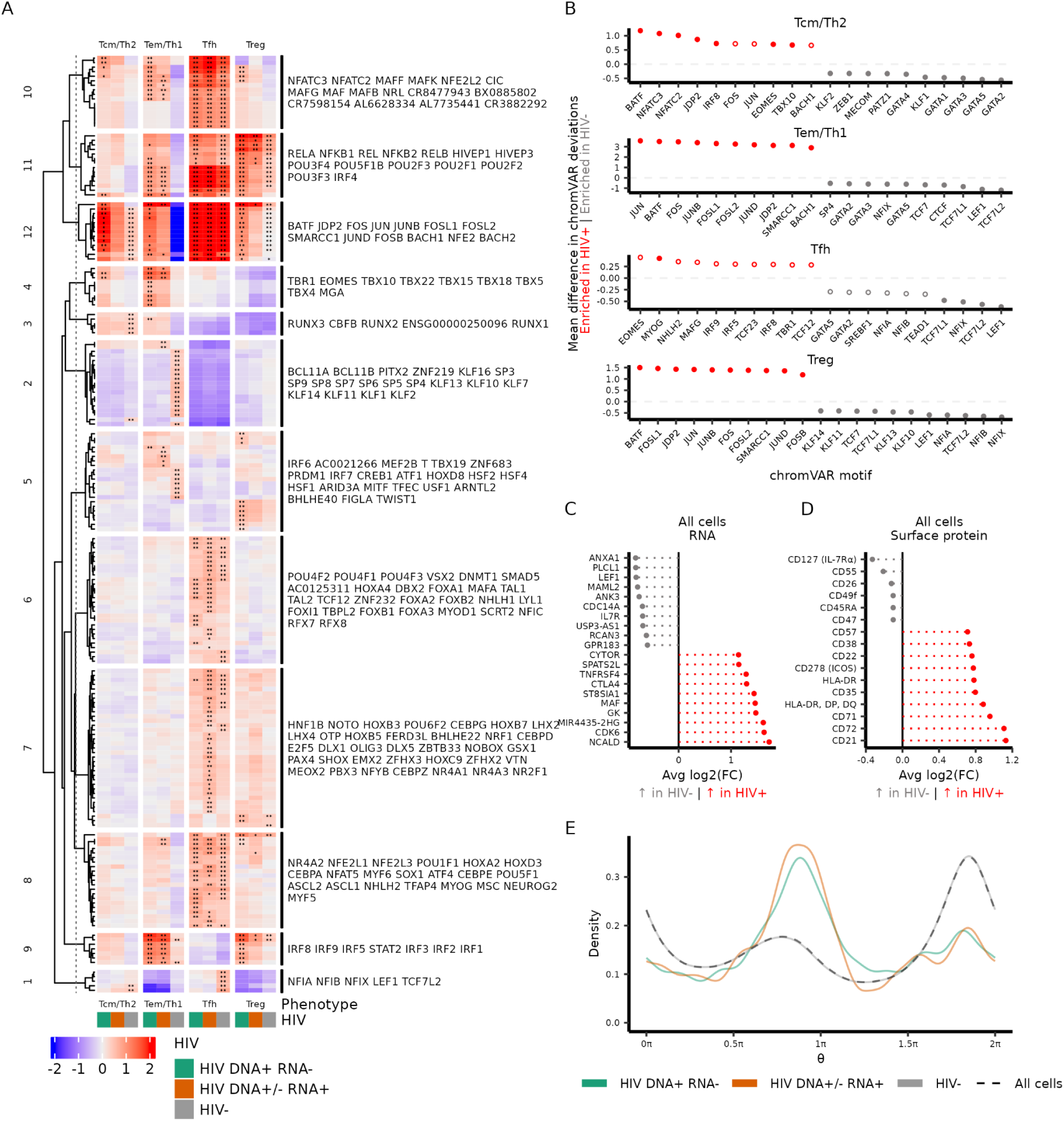
Heightened T-cell activation signatures in HIV+ cells during viremia. (A) Heatmap showing mean z deviation scores of transcription factor motifs from the CISBP database (displayed by row) in accessible DNA as calculated by chromVAR. Cells are grouped by HIV infection status and a binned annotation phenotype (displayed by column). Transcription factor motifs shown have been significantly enriched in a particular column over all other cells. Displayed motifs have been filtered (FDR < 0.05 and a mean difference > 0.5) to select for motifs that are found more in the accessible DNA of cells in that specific cluster (indicated by asterisks) relative to all other cells (getMarkerFeatures function in ArchR). Mean z deviation scores > 0 indicate a positive enrichment of a transcription factor motif. Rows were clustered using k-means clustering (with k = 12 to represent the number of columns). The motifs are labeled to the right of the heatmap in order from top to bottom for each clustered set of motifs. Asterisks indicate that the motif was significantly enriched in the specific group of cells. * = FDR < 0.05; ** = FDR < 0.01. (B) Differential chromVAR motifs were assessed in specific binned annotations between HIV+ (aggregate of HIV DNA+ RNA- and HIV DNA+/- RNA+) and HIV- cells. The top 10 motifs are shown in both directions of enrichment Solid dots represent a significant (FDR < 0.05) enrichment while open dots are not significant. (C) Differential gene expression between all cells in the dataset between HIV+ and HIV- cells. The top 10 significant genes, ranked by average log2(fold change), are shown in both directions (adjusted p value < 0.05). (D) Differential surface protein abundance between all cells in the dataset between HIV+ and HIV- cells. Up to the top 10 significant (adjusted p value < 0.05) surface proteins are shown in both directions. For both (C) and (D), only features were selected for differential comparison if there was at least 30% of cells with expression of the gene or protein in at least HIV+ or HIV- cells. Calculations were performed using the Wilcoxon Rank Sum test with Bonferroni-Hochberg correction. (E) Cell cycle position (θ where π is approximately G2M and 1.5π is middle of M stage) was estimated with tricycle^50^ to compare between HIV+ cells and HIV- cells. The distribution of cell cycle positions is shown based on the groups (shown by color). The black dashed line represents the distribution of all cells.

BATF and AP-1 family transcription factor motifs were also consistently enriched in HIV+ (combining HIV transcriptionally active and inactive cells) versus HIV- cells, when performing within-subset comparisons except for Tfh cells (**Figure 3B**). In the Tfh cells, we did not observe differential motif enrichment associated with activation when comparing HIV+ and HIV- cells, given the inherent activated state of germinal center Tfh cells as seen in our dataset (**Figure 3A**) and from other studies^42^. Similar to our previous studies^9^, HIV- cells also had significantly higher enrichment of motifs such as TCF7 and LEF1 (**Figures 3A-B**) that regulate T cell differentiation and activation^43^.

Progressing through the central dogma, we next investigated whether there were significant transcriptomic differences between all HIV+ and all HIV- cells in viremic LNs. Of note, HIV+ cells had increased expression of CDK6 and MAF transcripts (**Figure 3C**), both important for regulating cell cycle and HIV infection^44,45^. At the cell surface level (**Figure 3D**), HIV+ cells expressed more proteins associated with activation and proliferation such as HLA-DR and CD71^46,47^. Given the known relationship of HIV to the cell cycle^48,49^, we next investigated whether LN HIV+ cells in our study were in different phases of the cell cycle compared to LN HIV- cells. After inferring cell cycle position using the tricycle package^50^, we observed that HIV+ cells had a significantly higher distribution of cells in the G2/M phases compared to the G0/G1 phases relative to the distribution observed with HIV- cells (adj p = 4.4e-16; Kolmogorov-Smirnov test with Holm multiple test adjustment; **Figure 3E**). HIV DNA+/- RNA+ cells did not have a significantly different cell cycle distribution compared to HIV DNA+ RNA-cells (adj p = 0.24) suggesting a HIV-specific effect on cell cycling during viremia (**Figure 3E**).

We further examined whether differential expressions from our aggregated dataset were driven by Tfh cells. The differentially expressed genes between HIV+ Tfh cells and HIV- Tfh cells differed compared to the aggregated dataset but also contained genes related to cell cycling. HIV+ Tfh cells expressed higher levels of CYTOR and BIRC3 transcripts (**Supplemental Figure 4A**) which have further implications for HIV replication^51,52^. Surface protein analysis of HIV+ Tfh cells also had partial agreement with the aggregate analysis with higher levels of CD71 and HLA-DR, albeit at a smaller fold change (**Supplemental Figure 4B**). HIV- Tfh cells also exhibited a pronounced peak at the G2/M phase which mirrored most other HIV+ subsets, likely due to their overall heightened activation level, though HIV+ Tfh cells were still higher (**Supplemental Figure 4C**). Together, our data indicate that many HIV+ cells show activation profiles and are actively entering the cell cycle compared to their HIV- counterparts in the lymphoid compartment during viremia.

### HIV+ cells during ART are largely indistinguishable from HIV- cells within the same CD4 T cell subset

To investigate potential differences between HIV+ and HIV- cells during ART, we performed an unbiased differential gene module analysis at subset resolution by identifying genes that are enriched in HIV+ (regardless of HIV RNA due to lower numbers) or HIV- cells compared to all other cells in the dataset (**Figure 4A**). We chose transcriptomic analysis in lieu of epigenetic analysis as ATAC is inherently more restricted by smaller sample sizes^53^. The resultant gene modules were primarily driven by T cell subset differences rather than HIV infection. These included T cell activation genes (JUN, FOS; Module 12), found primarily in Tem cells; T cell differentiation and cell cycling (TOX, CDK6, CD200, TIGIT; Modules 1 and 5) in Tfh cells; and regulatory T cell (Treg) associated genes (IKZF2, S100A4; Module 7) in Treg cells (**Figure 4A**).

**Figure 4:**
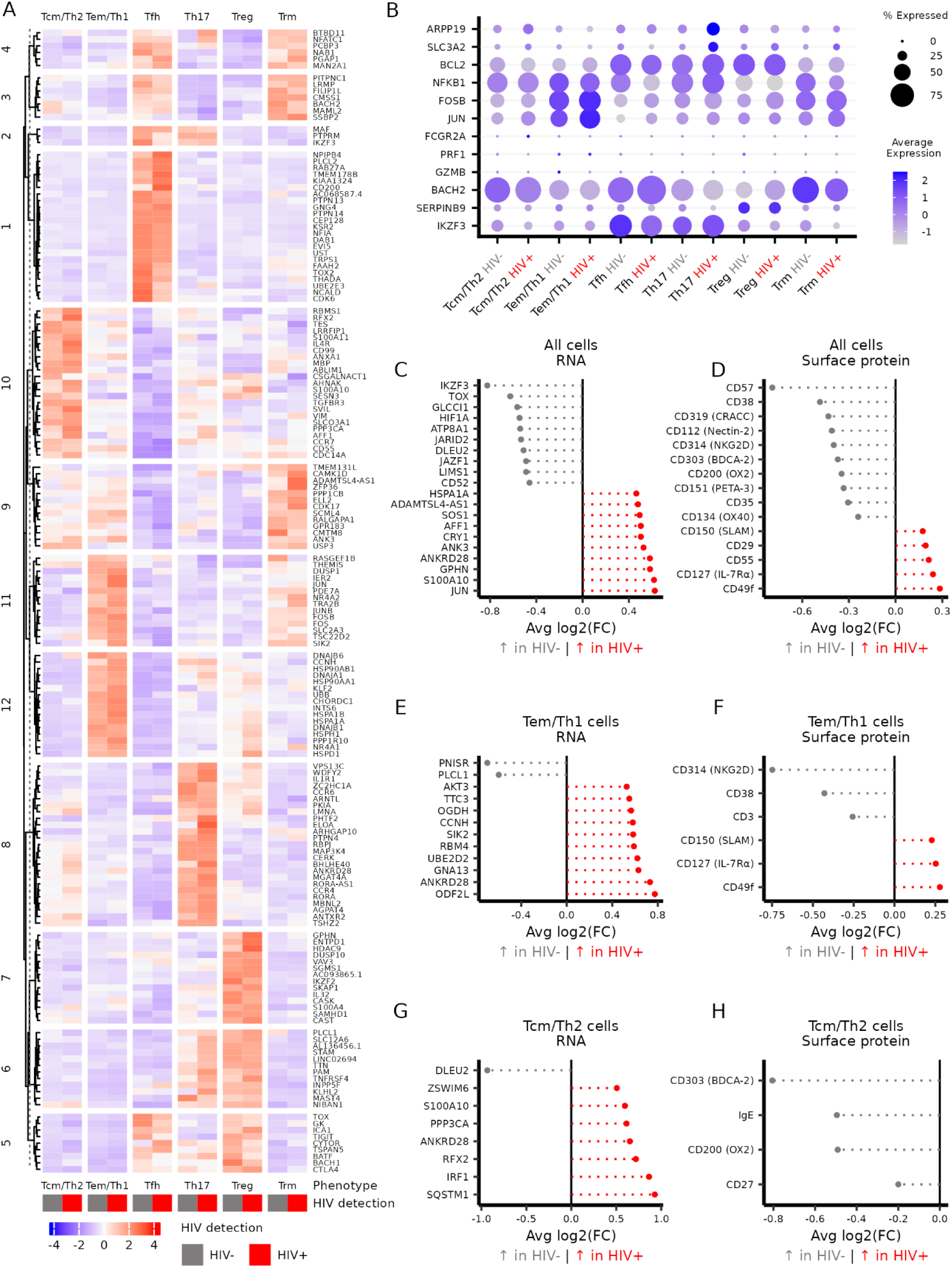
HIV+ cells during ART are largely indistinguishable from HIV- cells within the same CD4 subset. (A) Heatmap showing scaled (by row) gene expression of the top 25 (by average log2 fold change) genes that at least 30% of cells have expression of and were significantly enriched (adjusted p < 0.05) in a particular cell group (i.e. a column) over all other cells. Wilcoxon Rank Sum test with Bonferroni adjustment was used (as part of the FindAllMarkers function in Seurat). (B) Dot plot showing RNA expression of select genes that have been previously reported for HIV reservoir enrichment. (C and D) Differentially expressed genes and surface protein, respectively, are shown for all cells in the dataset between HIV+ and HIV- cells. (E and F) Differentially expressed genes and surface protein, respectively, are shown for Tem/Th1 cells between HIV+ and HIV- cells. (G and H) Differentially expressed genes and surface protein, respectively, are shown for Tcm/Th2 cells between HIV+ and HIV- cells. For (C) through (H), only features that have at least 30% expression and adjusted p value < 0.05 are shown from a Wilcoxon Rank sum test with Bonferroni-Hochberg adjustment (as calculated by the FindMarkers function in Seurat).

We next examined whether previously defined markers reported to enrich for HIV-infected cells in peripheral blood and gut^8–10,12,54,55^ could differentiate HIV+ cells from ART LN. We conducted differential gene expression of these specific markers within subset-matched HIV+ or HIV- cells in our dataset (**Figure 4B**). Of those genes with at least 25% of expression in either HIV+ or HIV- cells, we found no significant enrichment of any previously identified markers associated with HIV-infected CD4+ T cells beyond overall subset associations. For instance, SERPINB9 was more enriched in Tregs as a whole, but did not differ between HIV- and HIV+ cells.

Additional pathway enrichment ^56^ with UCell-based scoring^57^ found no significant enrichment between HIV- and HIV+ cells (**Supplemental Figure 5A**). Previous works have also suggested that HIV-infected cells during ART are resistant to cell death/apoptosis through a number of different pathways^8,12,54,58^. However, when comparing HIV+ vs HIV- cells from the four major infected subsets, we found no consistent pattern of differential transcript expression for pro-apoptotic and anti-apoptotic genes ^59^ across all HIV+ subsets (**Supplemental Figure 5B)**.

We subsequently compared all aggregated (i.e. not subset matched) HIV+ cells versus HIV-cells for differentially expressed transcripts (**Figure 4C**) and surface proteins (**Figure 4D**). We observed only subtly enriched transcripts in HIV+ cells with the top hit being JUN, while in contrast to a recent report^10^ IKZF3 (Aiolos) was enriched in HIV- cells. Subtle changes were also detected for surface protein expression with CD49f, CD127, CD55, CD29, and SLAM being increased on HIV+ cells, suggesting a heterogenous mixture of cells that differ from the patterns previously observed in peripheral blood^9^. HIV- cells had higher CD57 expression (**Figure 4D**), suggesting that HIV+ cells may be less enriched in GC-Tfh cells as GC-Tfh cells express CD57^60^. We did not observe the large changes in cell cycling genes that we had observed from the viremic samples. After inference of cell cycle state, we found that the distribution of cell cycle state did not significantly differ between HIV+ and HIV- cells (**Supplemental Figure 5B**) indicating that the cell cycle distortion in HIV+ cells was largely restricted to viremic infection.

Given the heterogeneous signal found in the aggregate comparison, we then assessed differential gene and protein expression between HIV+ and HIV- cells within the largest infected cell subsets Tem/Th1 (n = 132 infected cells) and Tcm/Th2 cells (n = 88 infected cells) (**Figures 4E-H**) . Overall, there were no strong differences between HIV+ and HIV- Tem/Th1 cells.

However, there was a subtle decrease in CD3 surface protein expression and increase in CD49f, CD127, and CD150 in Tem/Th1 HIV+ cells compared to Tem/Th1 HIV- cells (**Figure 4F**). In analysis of the HIV+ Tcm/Th2 cells, we also did not observe any major differences between HIV+ and HIV- cells at both transcript and surface protein expression (**Figures 4G-H**). ANKRD28 transcript, a gene known to be important for cell migration^61^, was increased at subtle levels in HIV+ compared to HIV- cells for the Tcm/Th2, Tem/Th1, and aggregated comparisons. Taken together, these data suggest that within LN during ART, HIV+ CD4 T cells are largely indistinguishable from subset-matched HIV- CD4 T cells at the transcriptomic and proteomic level.

### Specific depletion and infection of PRDM1+ Tfh cells during viremic HIV-1 infection

Given the propensity for Tfh cells to harbor HIV during viremic infection, we further explored the three different identified GC-Tfh clusters (**Figures 1C and 2)** to better understand potential mechanisms underlying their lack of contribution to the reservoir during ART. These three GC-Tfh clusters shared high locus accessibility and gene expression of the canonical Tfh markers *PDCD1* (PD-1) and *CXCR5* (**Figures 5A-B)**. However, they differed with regard to the key GC-Tfh transcription factor *BCL6*, in that GC-Tfh #1 and #2 clusters had high *BCL6* accessibility and transcription. In contrast, the third GC-Tfh cluster, PRDM1+ GC-Tfh, had comparatively lower *BCL6* accessibility and expression, higher locus accessibility and transcription for *PRDM1* (encoding the transcription factor BLIMP1), and higher IL-21 gene expression. While bearing some hallmarks of Tregs with increased locus accessibility and transcription of *TGFΒ* and *IL2RA*, the PRDM1+ GC-Tfh cluster did not express FOXP3 or IKZF2 (**Figures 5A-B, Supplemental Figure 2A-B)**, suggesting similarities to recently described non-canonical GC-Tfh subsets and distinction from T follicular regulatory cells (Tfrs) which express FOXP3^62,63^. The PRDM1+ GC-Tfh population was also unique in its increased locus accessibility and transcription of *CCR5* relative to all other LN CD4 T cell subsets (**Figures 5A-B, Supplemental Figure 2A-B**).

**Figure 5:**
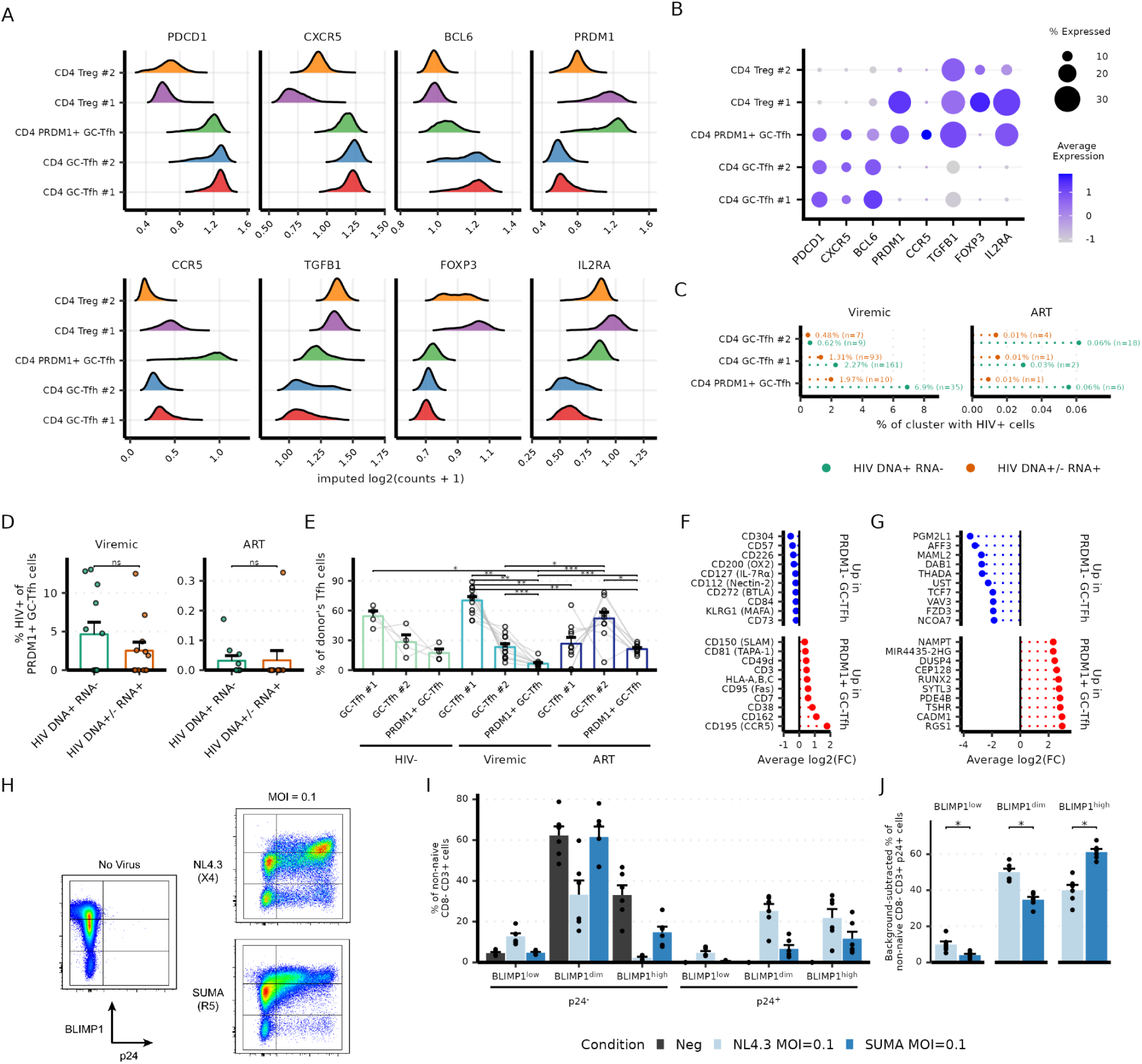
Specific depletion and infection of BLIMP1+ Tfh cells during viremic HIV-1 infection. (A) Imputed gene activity score from the ATAC modality (calculated by ArchR and imputed using MAGIC) for select Tfh and Treg markers. Values represent log2(normalized counts + 1). (B) Scaled average RNA expression for select Tfh and Treg markers. (C) Aggregate percentage of cells (irrespective of individual) within a cluster (separated by HIV infection stage) that has either HIV DNA+ RNA- (green) or HIV DNA+/- RNA+ (orange) cells. (D) Percentage of HIV+ cells within the PRDM1+ GC-Tfh cluster when stratified by individual. Statistical comparisons performed with Wilcoxon Rank Sum test and p-values were adjusted with Holm multiple test correction. (E) Distribution of donor’s Tfh cells (GC-Tfh #1, GC-Tfh #2, and PRDM1+ GC-Tfh) by HIV infection stage. Grey lines represent the same donor. Statistical comparisons performed with Wilcoxon Rank Sum test and p-values were adjusted with Holm multiple test correction. (F) Differential protein expression between all PRDM1- GC-Tfh cells (combination of GC-Tfh #1 and GC-Tfh #2) and PRDM1+ (GC-Tfh cells) in the aggregate dataset. The top 10 proteins (by average log2(fold change)) in either direction are shown. (G) Same as (F) but showing differential gene expression. For both (F) and (G), features that were tested required a minimum of 30% percent of cells that expressed the features in either cell subset. All features shown are significant (adjusted p-value < 0.05) by Wilcoxon Rank Sum test with Bonferroni-Hochberg multiple test correction. (H) Representative flow plots of tonsil organoid infection with either NL4.3 (X4 trophic virus) or SUMA (R5 trophic virus) at MOI 0.1. (I) Distribution of cells based on BLIMP1 levels and p24 positivity. Each dot represents a biological replicate and bars are colored by MOI and HIV. (J) Percentage of non-naive CD8-CD3+ p24+ cells within each gate (i.e. BLIMP1^high^) were background subtracted using the negative, uninfected control. Percentages relative to only p24^+^ cells were then calculated and shown. Statistical comparisons were performed with Wilcoxon Rank Sum Test followed by Bonferroni-Hochberg multiple test adjustment. * < 0.05

When assessing the percentage of infected cells within any given cluster, PRDM1+ GC-Tfh cells were the largest infected cell type amongst all memory CD4 T cell clusters with 8.8% of this cell population across viremic donors being HIV+ (**Figure 5C and Supplemental Figure 6A**). The high infection rate of CD4 PRDM1+ GC-Tfh cells was reflected in multiple viremic donors (**Figure 5D**) with several individuals having over a 10% infection rate in this specific subset. In addition, CD4 PRDM1+ GC-Tfh cells were highly depleted in viremic individuals (n = 507 out of 163061 cells; 0.31%) but recovered in ART-treated individuals (n = 10794 out of 294465 cells; 3.67%) (**Figures 5C, E**). The extent of depletion was quite pronounced given that the ratio of PRDM1+ Tfh cells to PRDM1- Tfh cells was at a 1:20.8 ratio during viremia compared to a 1:3.9 ratio after ART (**Figure 5C**).

To further examine how CD4 PRDM1+ GC-Tfh cells differed from the other GC-Tfh clusters, we assessed differential expression of surface proteins and transcripts between all PRDM1+ and PRDM1- GC-Tfh cells (**Figures 5F-G**). CCR5 and CD162 were the strongest differentially expressed surface proteins on PRDM1+ GC-Tfh cells (**Figure 5F**) - both of which have known roles as the coreceptor for HIV infection or for virion assembly/incorporation, respectively^64,65^.

These differences were also seen when comparing within HIV-, viremic, or ART-treated stages (**Supplemental Figure 6B**). The PRDM1+ GC-Tfh cells also had higher expression of CD38 and lower expression of CD127 (**Figure 5F**) as well as increased transcriptional expression of RUNX2 and CADM1 (**Figure 5G**).

Given the significant increase in CCR5 surface expression, we next assessed whether BLIMP1+ GC-Tfh cells were preferentially infected in vitro. Using purified CD4+ T-cells from people without HIV, we activated PBMC for 3 days, followed by infection with a R5-tropic HIV (SUMA strain) or X4-tropic HIV (NL4.3 strain) for 24 and 48 hours. In both R5 and X4 tropic strains, the vast majority of p24+ cells were BLIMP1+ and CCR5+ (**Supplemental Figures 7A-D**) after 24 and 48 hours post infection.

To avoid the need for in vitro T cell activation and instead have a direct look at the real susceptibility of different lymph node subsets, we next examined whether there would be preferential infection of BLIMP1+ cells in a tonsillar organoid model where BLIMP1 is already expressed by a subset of Tfh cells^62,66,67^ (**Supplemental Figure 7E**). We infected tonsil organoids with HIV SUMA or NL4.3 at a MOI of 0.1 for 3 days without exogenous activation. After combinatorial gating for BLIMP1 expression (in increasing order: BLIMP1^low^, BLIMP1^dim^, and BLIMP1^high^) and p24 expression (**Figure 5H**), we observed a large proportion of BLIMP1^high^ p24- cells in the negative control. As expected, this BLIMP1^high^ p24^-^ population was substantially reduced in infected conditions that corresponded with an increase in BLIMP1^high^ p24^+^ cells (**Figure 5I**). In most infection conditions, over 50% of the p24^+^ cells had BLIMP1^high^ expression (**Figure 5J**), highlighting the relationship between HIV infection/replication and BLIMP1. Together these data indicate that BLIMP1 expressing CD4+ T cells from lymphoid tissue are inherently susceptible to HIV infection and replication.

## Discussion

While there may be different paths towards a cure for HIV, some strategies may require the elimination or durable control of a persistent reservoir that is known to be pervasive and predominant in tissue environments at every infection stage and during ART. In light of this difficult context, a deeper understanding of the infected cells is critical for targeted clearance efforts in a therapeutic, safe, and cost-effective strategy given the wide spectrum of CD4+ T cells. Here, we have used a robust multimodal single-cell approach to create a comprehensive atlas of HIV infected cell identity, phenotype, and regulation from the LN of treated and untreated PWH. These data form a new foundation for our understanding of HIV tissue reservoirs that, in combination with recent efforts to more deeply understand HIV reservoirs in the gut^55,68,69^, have major implications in the global curative efforts for HIV.

The concept that follicular T cells, and later, Tfh cells, dominate the LN HIV reservoir has been a cornerstone of HIV reservoir biology and cure efforts for over 30 years^13,15,70,71^. While our data do not dispute that Tfh cells are an important part of the HIV reservoir, one of our major findings is that HIV LN reservoirs are not predominantly restricted to B cell follicles during both viremia and ART. In the absence of therapy, GC-Tfh cells were indeed one of the top contributors to the pool of infected cells (for either HIV DNA+ or RNA+), but were not the majority of infected cells. This is in agreement with previous reports where Tfh cells are proportionally more infected, but extra-follicular cells in total constituted a larger portion of the HIV LN reservoir^16^. It has long been assumed that Tfh cells also constitute the predominant LN reservoir during ART ^72^. We instead found that Tfh cells contribute only a fraction of the total infected cell reservoir, and various non-follicular Tem subsets dominated the ART LN reservoir. While it remains possible that Tfh may make a larger contribution in more activated LN such as those draining the mucosa and gut^23,73^ during ART, we did not observe a pattern between the overall frequency of Tfh cells between cervical or inguinal LN.

While Tfh cells were not the majority of infected cells, their importance in HIV immunology cannot be understated and is supported by our major finding of the selective depletion and infection of a PRDM1+ Tfh subset during viremia that recovers during ART. *PRDM1* encodes a transcription factor known as BLIMP1, which has a multifaceted role in CD4 T cell development. During GC-Tfh development, BCL6 is the main transcription factor responsible for driving the differentiation program and directly represses *PRDM1* transcription^74^. However, GC-Tfh cells are heterogeneous and recent publications in mice and humans have identified GC-Tfh derived subsets that selectively induce antibody secreting B cell formation or lead to the formation of inducible T follicular regulatory (iTfr) cells for regulating the development of antibody secreting cells^62,63^. One such subpopulation of GC-Tfh cells can re-express BLIMP1 in the progression towards a non-canonical Tfh cell and an iTfr phenotype^62^. BLIMP1+ Tfh cells are known to be important for providing B cell help^62,63^. In mice, selective knockout of *PRDM1* in CD4 T cells results in the loss of antibody-secreting plasma cells^63^. The depletion or disruption of BLIMP1+ Tfh cells during viremia could explain the observed loss of HIV-specific antibody secreting cells during chronic infection relative to early infection ^75^. We also observed that of the 3 GC-Tfh clusters, the PRDM1+ GC-Tfh cells expressed the most IL-21 transcripts - encoding a cytokine critical for GC B cell affinity maturation and transition to antibody secreting B cells^76^. Previous studies have demonstrated that the absence of IL-21 underlies impaired B cell responses in untreated HIV infection^77^, which may be explained by the selective depletion of the PRDM1+ GC-Tfh population in viremic infection. Given the propensity for these cells to be infected, it is also highly relevant to consider PRDM1+ GC-Tfh cells as a flashpoint for recrudescent viral replication if ART is interrupted. Although they do not appear to harbor a meaningful reservoir during ART, they would perhaps be the most susceptible CD4 population in the LN for new rounds of replication.

We were unable to corroborate recent reports highlighting purported associations between specific protein signatures and/or transcriptional programs and the HIV infected cell reservoir in blood^8–10,12,54^ or gut^55^. This could reflect a tissue-restricted phenomenon, where specific phenotypes of cells, including the HIV+ cells, more directly reflect their environment (i.e. resident memory T cells in gut or effector/further differentiated cells in peripheral blood). While we found no singular marker or set of markers unique to HIV+ cells during viremia or ART treatment in LN, we were able to ascertain modest signatures of the HIV+ cell with our trimodal resolution. While overall similar to their HIV- counterparts, HIV-infected cells from viremic LN showed enrichment for several epigenetic modules. Throughout the different CD4+ T cell subsets, HIV+ cells had increased signatures related to AP-1 and NF-κB transcription factors. As these transcription factors are positive regulators of T cell proliferation and activation^78^ and bind in the proviral LTR^79–81^, the chromatin environment of HIV+ cells in our dataset appeared to be primed for HIV replication and transcription during viremia. Furthermore, the increased enrichment of interferon (IFN) related transcription factor motifs in different subsets of HIV+ cells validated their infected states given that HIV is known to trigger a cGAS-dependent type I IFN response in CD4+ T-cells^39^. In contrast with the signatures described during viremic infection, the HIV+ and HIV- cells during ART were largely similar to each other when compared within memory/differentiation subsets. We did find a large proportion of HIV+ cells during ART in subsets expressing *NR4A1* and/or *NR4A2* transcripts. *NR4A1* (Nur77) is used to identify very recent activation through the T cell receptor in transgenic mouse models^82^ and human T cells^83^. It is unclear if these cells have been recently activated besides *NR4A1*/*NR4A2* RNA expression as they also express other activation markers such as *JUN* and *FOS*. Overall, we could not find a defining signature that could solely describe the HIV+ cells, highlighting the challenge of specifically targeting the HIV reservoir in the tissue environment.

From our observations of differential transcript and protein expressions of cell cycling related genes, HIV+ cells were also found to be more skewed towards a G2/M phase compared to HIV-cells, with the exception of HIV- Tfh cells. The HIV replication cycle and host cell cycle are intertwined in many ways, including the importance of cyclin T1 for Tat-mediated transcription^84^ and the disruptive impact of viral Vpr on the cell cycle^48^. While it is unclear from our dataset whether HIV+ cells are actively dividing or arrested near the G2/M phase, HIV infection nevertheless has a profound impact on the cell cycle that may be useful for potential targeting. For instance, duramycin conjugated micelles selectively deliver anti-cancer drugs to cells at the G2/M phase^85^. However, the differential cell cycle related genes did not extend to ART-treated PWH, potentially suggesting that 1) the surviving HIV+ cells may be remodeled by ART or resolution of inflammation and/or 2) the HIV+ cells that survive to establish a reservoir after ART are different compared to infected cells that do not survive.

Our study has a number of limitations. As with all single-cell assays that use ATAC based detection, our detection of infected cells is inherently limited to cells with accessible proviruses which limit our ability to properly discern the latent reservoir. The infected cells that we do detect are still relevant to curative efforts as they still contribute to a significant part of the reservoir.

Furthermore, accessible DNA does not necessarily equate to a non-latent cell, as there are multiple routes to latency including transcriptional impediments^86^. Given the known ∼50% efficiency of ATAC based procedures^87^, it is not surprising that we and others^10,55^ would detect RNA+ cells without detectable viral DNA. As such, the DNA+ reservoir is likely larger than we can detect, due both to the efficiency of ATAC and the potential for heterochromatic provirus.

Another limitation is the inability to discern between intact and defective proviruses. It is likely that the majority of detected proviruses in infected cells are defective, but are still of importance given that the defective proviruses have been shown to still produce transcripts and proteins^88,89^ that would be of relevance to an immune response. Additional technical advances are urgently needed for an unbiased high-throughput and single-cell method to acquire complete proviral sequence along with multimodal phenotyping. The rarity of infected cells during ART also restricted the extent of analyses that were possible. Despite this, we were able to perform differential analyses between HIV+ and HIV- cells. It is also unclear whether the cervical and inguinal LNs are representative of every LN in the body, and the LN HIV+ T cell reservoir could have a different composition across LNs even within an individual depending on location and local immune reactivity. Addressing such issues is logistically challenging, but recent efforts to obtain multiple LN tissues from HIV+ organ donor programs such as Last Gift^90,91^ will help to address this critical question.

Our work has major implications for cure efforts and strategies that solely focus on reservoir clearance from LN B cell follicles during ART. Indeed, neither efforts to engineer CXCR5+ CAR T cells^92–94^ or alter the follicular environment^94–96^ in the SIV rhesus macaque model have been successful for reservoir clearance. Our findings instead highlight the need for combinatorial strategies that can target both follicular and non-follicular cells to comprehensively address the lymphoid HIV reservoir and further demonstrate the rationale for multimodal profiling to uncover new immune dynamics of HIV infection.

## Supporting information

Main Table 1

## Acknowledgements

We thank the individuals who donated samples to our study. We also thank members of the Betts Lab for their assistance. We would like to acknowledge assistance provided by various cores: the Penn Cytomics and Cell Sorting Shared Resource Laboratory at the University of Pennsylvania (RRID:SCR_022376); the Penn Genomics and Sequencing Core (RRID:SCR_022382), and the Virus & Reservoirs Technology Core at the Penn Center for AIDS Research. The authors thank Max Eldabbas, Emileigh Maddox, Tanishk Sinha, and Jiayi Shu of the Human Immunology Core (HIC) (RRID:SCR_022380) at the Perelman School of Medicine at the University of Pennsylvania for assistance with apheresis collection. Penn Cytomics is partially supported by the Abramson Cancer Center NCI Grant (P30 016520). The HIC is supported in part by NIH P30 AI045008 and P30 CA016520.

## Author contributions statement

VHW, JMLN, MRB conceptualized experiments. MBP and JMLN performed flow and sorting experiments. JMLN and VHW performed scDOGMAseq experiments and analysis. PMdRE, FTR, MGN, GS, YALV, GRT, and SAR recruited and organized sample collection and cell preparations used for scDOGMAseq. JMLN performed in vitro infection experiments. SR, NR, and JMLN organized and performed tonsil organoid infection experiments. JAR and KJB provided viral stocks used for in vitro infection experiments. WLB, MC, and JDE performed IHC and DNAscope experiments. MHS and OSS recruited and provided tissue samples for DNAscope validation experiments. VHW performed the bioinformatic analysis. VHW, JMLN, and MRB annotated the clusters. VHW, JMLN, and MRB wrote the manuscript.

## Methods

### Study approval

This study complied with all relevant ethical regulations and the principles of the Declaration of Helsinki. People without HIV (n = 4) and people with HIV (n = 22) were originally recruited at the Centro de Investigación en Enfermedades Infecciosas of the Instituto Nacional de Enfermedades Respiratorias “Ismael Cosío Villegas” (CIENI–INER; Mexico City, Mexico). All participants provided written informed consent for lymph node tissue donation under protocols B03-16 and B33-10, approved by the Ethics Committee and the Ethics in Research Committee of the INER and by the Institutional Review Board of the University of Pennsylvania. The use of these samples for the present study was additionally approved by the Research Committee and the Ethics in Research Committee of the Instituto Nacional de Enfermedades Respiratorias “Ismael Cosío Villegas” (protocol C71-18). All participants were provided with the results of all clinical tests performed, including tissue biopsy reports, plasma viral load measurements, and lymphocyte counts.

### scDOGMAseq cell processing and sorting

Inguinal and cervical LN were obtained from study participants and processed into single-cell suspensions, then cryopreserved as previously described^97,98^. Frozen lymph node mononuclear cells (LNMC) were briefly thawed at 37°C, washed, resuspended at 2 million cells/mL in complete R10 medium (RPMI + 10% FBS + 1% penicillin/streptomycin + 2 mM L-glutamine) with 10U/mL DNAse I (Roche Life Sciences), and rested for 2 hours in a humidified incubator at 37°C + 5% CO_2_. Following the 2 hour rest, samples were counted and separated for phenotype assessment or sorting. For sorting, cells were washed with PBS and prestained for the chemokine receptor CCR7 for 15 minutes in the incubator at 37°C + 5% CO_2_. All the following incubations were performed at room temperature in the dark. Cells were stained with viability dye for 10 minutes (Live/Dead Fixable Aqua, Invitrogen), followed by the extracellular antibody cocktail and each donor sample was stained with a unique hashing antibody (HTO, TotalSeq-A Hashing, BioLegend) for 20 minutes. After staining, cells were washed with PBS three times at 500 x g for 5 minutes to remove residual HTO antibody. Following the wash, cells were resuspended in PBS+1% BSA and set on ice.

Immediately prior to sorting, cells were pooled together and filtered through a 35um nylon mesh filter top tube to avoid cell clumping. All samples were sorted on a FACS Aria II SORP (BD) inside a biosafety cabinet for biohazardous samples following proper EHRS standards. The gating strategy is shown in supplemental figure 1B. Cells were sorted into PBS+1% BSA at 4°C and placed on ice for downstream analysis in scDOGMAseq.

The following antibodies were used for cell sorting: CCR7 APC-Cy7 (clone G043H7), CD16 PE-Cy5 (clone 3G8), and CD19 PE-Cy5 (clone HIB19) were acquired from Biolegend. CD3 APC-R700 (clone UCHT1), and CD45RA PE-CF594 (clone 5H9) were acquired from BD. CD14 PE-Cy5 (clone 61D3), and CD8 PE Cy5.5 (clone RPA-T8) were acquired from eBioscience.

### LNMC staining for phenotype assessment

After splitting the sample between phenotype assessment and sorting, cells were prestained for the chemokine receptor CCR7 and CXCR5 for 10 minutes at 37°C + 5% CO_2_. All following incubations were performed at room temperature in the dark. Cells were stained for viability dye for 10 minutes (Live/Dead Fixable Aqua,

Invitrogen), followed by surface markers staining (20 minutes). Cells were washed and fixed/permeabilized using the FoxP3 Transcription Factor Buffer Kit (eBioscience), following the manufacturer’s instructions. Intracellular staining was performed by adding the antibody cocktail prepared in a 1X permwash buffer and incubated for 1 hour at 37°C. Stained cells were washed and fixed in PBS containing 1% paraformaldehyde (Sigma-Aldrich), and stored at 4°C in the dark until acquisition. All flow cytometry data were collected on a BD FACSymphony A5 cytometer (BD Biosciences) and data were analyzed using FlowJo™ v10.8.1 Software (BD Life Sciences).

The following antibodies were used for phenotype assessment: CCR7 APC-Cy7 (clone G043H7), PD1 BV421 (clone EH12.2H7), CD14 BV510 (clone M5E2), CD19 BV510 (clone HIB19), CD27 BV570 (clone O323), CD38 BV711 (clone HIT2), CD69 PE-Cy5 (clone FN50) and Granzyme K PE (clone GM26E7) were acquired from Biolegend. CXCR5 BV750 (clone RF8B2), CD39 BB700 (clone TU66), CD4 BB790 (clone SK3), CD8 BUV496 (clone RPA-T8), HLA-DR BUV615 (clone 3G8), CD95 BUV737 (clone DX2), TIGIT BV605 (clone 741182), KI67 BUV395 (clone B56), Granzyme B AF700 (clone GB11), CD3 BUV805 (clone UCHT1) and CD45RA PE-CF594 (clone HI100) were acquired from BD. Tox APC (clone REA473) was acquired from Miltenyi Biotech. TCF1 AF488 (clone C63D9) was acquired from CST. Tbet PE-Cy7 (clone 4B10) was acquired from eBioscience.

### scDOGMAseq library preparation and sequencing

Buffer and cell preparation were performed as previously published following the LLL permeabilization protocol^24^. All subsequent cell preparation and antibody staining steps, including centrifugation, were performed at 10°C or on ice unless specified. 5 x 10^5^ - 1 x 10^6^ sorted CD3+, CD8- non-naive T cells were pelleted at 500 x g for 5 minutes. Cell pellet was resuspended in 22.5uL of Staining Buffer (PBS, 1% BSA, 0.2U/uL RNAse Inhibitor (Roche)) and incubated with 2.5uL of TruStain FcX (Biolegend) for 10 minutes. One vial of TotalSeq-A Human Universal Cocktail, v1.0 (Biolegend) was resuspended according to the Biolegend protocol with Staining Buffer. 25uL of prepared antibody cocktail was added to the cells and incubated for 30 minutes. Cells were washed three times with 1 mL of Staining Buffer at 400 x g for 5 minutes. After the final wash cells were resuspended in 450uL of PBS.

Following the LLL permeabilization protocol cells were fixed with 0.1% paraformaldehyde for 10 minutes at room temperature and quenched with 0.125M final glycine solution. Fixed cells were washed two times with Staining Buffer at 400 x g for 5 minutes. Cells were lysed with 100uL of LLL lysis buffer (10 mM Tris-HCl pH 7.4, 10 mM NaCl, 3 mM MgCl_2_, 0.1% NP40, 1% BSA, 1 mM DTT, 2 U/μl RNAse inhibitor) for 3 minutes. Cells were washed with 1mL of cold LLL wash buffer (10 mM Tris-HCl pH 7.4, 10 mM NaCl, 3 mM MgCl_2_, 1% BSA, 1 mM DTT, 1 U/μl RNAse inhibitor) and the tube inverted before pelleting the cells at 500 x g for 5 minutes. Cell pellet was resuspended with 10X Genomics Multiome 1x Nuclei Buffer, and complete cell permeabilization was confirmed using Trypan Blue staining. If aggregates were detected in the cell count, cells were filtered using a 40um FlowMi strainer (Sigma) and re-counted. Cells were diluted as needed according to the 10X Chromium Next GEM Single Cell Multiome ATAC + Gene Expression protocol (Version CG000338 RevF).

Samples were processed using the Chromium X platform (10X Genomics). Single-cell sequencing library preparations for the ATAC and GEX libraries were made following the 10X Genomics User Guide Chromium Next GEM Single Cell Multiome ATAC + Gene Expression protocol (Version CG000338 RevF), with the following modifications as previously published^24^ and outlined below.

- 10X protocol Step 4.1, during the preamplification PCR spike in 1 μL of 0.2 μM ADT/HTO additive primer (ADT additive primer sequence: 5’CCTTGGCACCCGAGAATT*C*C, HTOv2 additive primer sequence:5’GTGACTGGAGTTCAGACGTGTGCTCTTCCGAT*C*T) was added to each individual sample preparation.

- 10X protocol Step 4.3k, during the SPRI clean-up, elute the pre-amplification sample in 100uL of EB buffer. Use 25uL of pre-amplification sample +15uL of EB (Qiagen) as input for the ATAC library indexing, and use 35uL of pre-amplification sample as input for the cDNA amplification reaction. Use 5uL of pre-amplification sample as input for the ADT and HTO libraries.

ADT/HTO dual index single-cell sequencing library preparation followed the antibody manufacturer’s protocol (BioLegend). Briefly, 5uL of the pre-amplification sample was mixed with 40uL of H_2_0. The antibody-derived oligomers were amplified and indexed with 50 uL of KAPA HiFi HotStart ReadyMix (KAPA Biosystems) and 5 uL of forward/reverse indexing primers, outlined below (BioLegend).

The PCR amplification settings for the ADT modality were as follows:

● 95 °C for 3 minutes,
● 14–15 cycles of:

○ 95 °C for 20 seconds,

○ 60 °C for 30 seconds,

○ 72 °C for 20 seconds,

● 72 °C for 5 minutes,
● Hold at 4 °C.

The PCR amplification settings for the HTO modality were as follows:

● 95 °C for 3 minutes,
● 13–15 cycles of:

○ 95 °C for 20 seconds,

○ 64 °C for 30 seconds,

○ 72 °C for 20 seconds,

● 72 °C for 5 minutes,
● Hold at 4 °C.

Post-amplification, ADT/HTO libraries were purified with 120 uL SPRIselect (Beckman) cleanup and eluted with 30 uL of EB buffer. All final libraries were quantified using a Qubit dsDNA HS Assay Kit (Invitrogen) and analyzed using a High Sensitivity D1000 DNA tape (Agilent) on a TapeStation D4200 (Agilent).

ADT and HTO Indexing Primers

- Dual Index ADT i5/i7 Primer Pairs (DI_ADTx) (for ADT amplification) [DI_ADT1 i5] - 5’AATGATACGGCGACCACCGAGATCTACAC-[10 bp index]-ACACTCTTTCCCTACACGACGC*T*C

[DI_ADT1 i7] - 5’CAAGCAGAAGACGGCATACGAGAT-[10 bp index]-GTGACTGGAGTTCCTTGGCACCCGAGAATTC*C*A

- Dual Index HTO i5/i7 primer pairs (DI_HTOx) (for HTO amplification)

[DI_HTO1 i5] - 5’AATGATACGGCGACCACCGAGATCTACAC-[10 bp index]-ACACTCTTTCCCTACACGACGC*T*C

[DI_HTO1 i7] - 5’CAAGCAGAAGACGGCATACGAGAT-[10 bp index]GTGACTGGAGTTCAGACGTGTGCTCTTCCGAT*C*T

Sequencing was performed on the NovaSeq 6000 platform (Illumina) with a target of at least 25,000 reads per cell for ATAC libraries, 25,000 reads per cell for RNA libraries, 10,000 reads per cell for ADT libraries, and 1,000 reads per cell for HTO libraries. Libraries were denatured and diluted according to the Illumina NovaSeq 6000 Protocol A: Pool and Denature Libraries for Sequencing (Standard Loading, Document #1000000106351 v03). Additionally, 1-2% PhiX was added to the library pool for quality control and enhanced diversity (1% for ATAC libraries, and 2% for GEX, ADT and HTO libraries). Sequencing geometries followed the cycle length outlined in the User guide 10X Chromium Next GEM Single Cell Multiome ATAC + Gene Expression protocol (Version CG000338 RevF).

*scDOGMAseq analysis*: Raw BCL files from sequencing for ADT/HTO modalities were demultiplexed using bcl2fastq2 (Illumina) or cellranger-arc (10X Genomics) mkfastq. ATAC fastq files were filtered for index hopping using index-hopping-filter (10X Genomics). ATAC and GEX fastq files were then aligned to a chimeric hg38-HIV (HXB2 sequence) genome and count matrices were generated using cellranger-arc count. ADT and HTO fastq files were aligned to the TotalSeqA cocktail catalog using kallisto and bustools^99^ which includes barcode correction and counting to create cell by feature matrices. A Snakemake pipeline for kallisto and bustools is found here: https://github.com/betts-lab/scc-proc. After alignment, barcodes associated with multiplets were detected using AMULET^100^ (false discovery rate (FDR)-corrected (Bejamini–Hochberg method) P value > 0.01 is denoted as a multiplet; default parameters).

Fragment files were loaded into ArchR^53^ and filtered for singlet barcodes that were identified with AMULET. The filtered feature barcode matrices for the RNA modality (output of cellranger-arc count) were loaded into ArchR as well. Only cell barcodes with both valid ATAC and RNA counts were kept for additional processing. Cells were further filtered by a stringent criteria of quality control metrics: transcription start site enrichment score >= 8, log(number of unique ATAC fragments) >= 3.6, number of detected genes with RNA modality > 500, number of detected genes with RNA modality < 6000, and percentage of mitochondrial-associated RNA counts < 10%. The ADT and HTO modalities were then loaded into a Seurat^101^ object after filtering for QC-passed barcodes from the RNA and ATAC modalities. HTO counts were normalized and unhashed using Seurat (function MULTIseqDemux with automatic threshold). Unhashed and singlet cells were retained for all modalities for downstream analysis. RNA counts were log normalized and ADT counts were centered log ratio transformed with a scale factor of 10,000. Module scores were generated for gene sets associated with B and APC/NK cells to remove contaminating cells. Iterative latent semantic indexing (LSI) was performed using ArchR on the ATAC and RNA modalities to generate a reduced dimensional space. Batch effects were corrected using donor and chip (i.e. experimental run) information using Harmony^102^. Both ATAC and RNA LSI spaces were used to generate clusters with a combined Seurat and ArchR approach. All analysis code is available at https://github.com/betts-lab/hiv-ln-dogma. FASTQ and processed files are deposited in GEO under accession number GSE313072.

### Identification of HIV+ cells

Using the BAM files from the cellranger-arc output, these were namesorted and ran through hiv-haystack^9^ to compile a list of cell barcodes with proviral reads. Valid HIV RNA reads were obtained by extracting all BAM records with mapping to the HXB2 viral genome and using the same thresholds that were previously published by a separate group^10^ to control for background noise and index hopping. A cell was considered HIV RNA+ if it had two or more viral RNA reads with different UMIs associated with its cell barcode or if it had four or more (PCR/optical) copies of a single viral RNA read.

### CD4+ T cell infection

Enriched CD4+T cells were obtained from an HIV-negative donor apheresis from the Human Immunology Core at the University of Pennsylvania. 3 - 5 x 10^6^ cells were set aside and resuspended in 3mL of complete R10 for a no stimulation control. The remaining cells were resuspended at a concentration of 2 x10^6^ cells/mL in complete R10 and activated with anti-CD3 (1 μg per mL; Bio-Rad), recombinant IL-2 (100 U per mL; Sigma Millipore), and anti-CD28 + anti-CD49d (1.5 μg per mL; BD) for 3 days in a humidified incubator at 37°C+5% CO_2_. Activated CD4+ T cells were resuspended in 1 mL of complete R10 media. HIV-1 virus stock (strain SUMA; provided by the University of Pennsylvania CFAR Virus and Reservoirs Core, and strain NL4.3; provided by the Bar Lab University of Pennsylvania) was thawed briefly in a 37°C water bath; 0, and 0.1 MOI equivalent of viral stock was added to the suspension and mixed by pipetting. Cells were then infected by spinoculation with polybrene for 45 minutes at 400 x g. Cells were rested in a humidified incubator at 37°C+5% CO_2_ for 1 hour (cap loosened to promote gas exchange). After resting, cells were washed with complete R10. Cells were resuspended at 2 x10^6^ cells per mL in complete R10 and left to rest in a 6 well plate. 24 hours after infection ½ of the cells were collected for staining and the volume replaced with complete R10. 48 hours after infection the total cells in the well were collected for staining.

Samples were prestained for the chemokine receptors CCR5, CCR7 and CXCR5 for 10 min at 37°C + 5% CO_2_. All following incubations were performed at room temperature in the dark. Cells were stained for viability dye for 10 minutes (Live/Dead Fixable Aqua, Invitrogen), followed by the extracellular antibody cocktail for 20 minutes. Cells were washed and fixed/permeabilized using the FoxP3 Transcription Factor Buffer Kit (eBioscience), following the manufacturer’s instructions. Intracellular staining was performed by adding the antibody cocktail prepared in a 1X permwash buffer for 1 hour. Stained cells were washed and fixed in PBS containing 1% paraformaldehyde (Sigma-Aldrich), and stored at 4°C in the dark until acquisition. All flow cytometry data were collected on a BD FACSymphony A5 cytometer (BD Biosciences) and data were analyzed using FlowJo™ v10.8.1 Software (BD Life Sciences).

The following antibodies were used for CD4+ T cell analysis: CCR5 APC (clone J418F1), CCR7 APC-Cy7 (clone G043H7), CD19 BV786 (clone HIB19), CD14 BV510 (clone M5E2), CD8 BV570 (clone RPA-T8), CD27 BV650 (clone O323), PD1 BV750 (clone EH12.2H7), and CD69 PE-Cy5 (Clone FN50) were obtained from Biolegend. CXCR5 BUV661 (clone RFB2), CD4 BB790 (clone SK3), CD45RA PE-CF594 (cloneHI100), CD38 BUV395 (clone HIT2), CD25 BUV496 (clone 2A3), ICOS BUV615 (clone DX29), CD71 BV605 (clone M-A712), CD3 BUV805 (clone UCHT1), BLIMP1 BV421 (clone 6D3), BCL6 RB705 (clone K112-91), and KI67 AF700 (clone B56) were obtained from BD. FoxP3 PE-Cy7 (clone 236A/E7) was obtained from Invitrogen. HIV-1 p24 RD1 (clone KC57) and HIV-1 p24 FITC (clone KC57) were obtained from Beckman Coulter.

### Tonsil organoid infection

As previously described ^62^, tonsils were acquired after tonsillectomies from deidentified immune-competent children that were ordered to address airway obstruction or recurrent tonsillitis. As such, the tonsils were designated as discarded surgical waste, and therefore this protocol was not classified as human subjects research by the Children’s Hospital of Philadelphia Institutional Review Board. A single-cell suspension of tonsillar mononuclear cells was created by mechanical disruption (tonsils were minced and pressed through a 70-μm cell screen) followed by Ficoll-Paque PLUS density gradient centrifugation (GE Healthcare Life Sciences). Mononuclear cells were counted and resuspended in organoid media [RPMI 1640 with L-glutamine, 10% FBS, 2 mM glutamine, 1× penicillin-streptomycin, 1 mM sodium pyruvate, 1× minimum essential medium nonessential amino acids (NEAA), and 10 mM HEPES buffer].

In a 15 ml conical tube, 6 × 10^6^ mononuclear cells were resuspended in 1 mL of organoid media. HIV-1 virus stock (strain SUMA; provided by the University of Pennsylvania CFAR Virus and Reservoirs Core, and strain NL4.3; provided by the Bar Lab University of Pennsylvania) was thawed briefly in a 37°C water bath; 0, and 0.1 MOI equivalent of viral stock was added to the suspension and mixed by pipetting. Cells were then infected by spinoculation with 8ug of polybrene for 45 min at 400 x g . Cells were rested in a humidified incubator at 37°C+5% CO_2_ for 1 hour (cap loosened to promote gas exchange). After resting, cells were washed and resuspended in 100 μL of organoid media. The total cell volume was transferred to permeable transwells (0.4-μm pore, 12-mm diameter; Millipore). Transwells were then inserted into standard 12-well polystyrene plates containing 1 ml of additional organoid media supplemented with recombinant human B cell activating factor (rBAFF) (1 μg/ml); BioLegend] and placed in an incubator at 37°C + 5% CO_2_ for up to 7 days. Media outside of the transwell was refreshed after 3 days without disrupting the organoids in the transwell.

3 days after spinfection organoids were dispersed via pipetting and washed with PBS. Samples were prestained for the chemokine receptors CCR5, CCR7 and CXCR5 for 10 min at 37°C + 5% CO_2_. All following incubations were performed at room temperature in the dark. Cells were stained for viability dye for 10 minutes (Live/Dead Fixable Aqua, Invitrogen), followed by the extracellular antibody cocktail for 20 minutes. Cells were washed and fixed/permeabilized using the FoxP3 Transcription Factor Buffer Kit (eBioscience), following the manufacturer’s instructions. Intracellular staining was performed by adding the antibody cocktail prepared in a 1X permwash buffer for 1 hour. Stained cells were washed and fixed in PBS containing 1% paraformaldehyde (Sigma-Aldrich), and stored at 4°C in the dark until acquisition. Immediately prior to acquisition, samples were filtered through a 35um nylon mesh filter top tube to remove cell clumps. All flow cytometry data were collected on a BD FACSymphony A5 cytometer (BD Biosciences) and data were analyzed using FlowJo™ v10.8.1 Software (BD Life Sciences).

The following antibodies were used for tonsil organoid analysis: CCR5 PE-Cy7 (clone J418F1), CCR7 APC-Cy7 (clone G043H7), CD19 BV786 (clone HIB19), CD14 BV510 (clone M5E2), CD8 BV570 (clone RPA-T8), CD27 BV650 (clone O323), PD1 BV750 (clone EH12.2H7), and CD69

PE-Cy5 (Clone FN50) were obtained from Biolegend. CXCR5 AF647 (clone RFB2), CD127 BV711 (clone HIL-7R-M21), CD4 BB790 (clone SK3), CD45RA PE-CF594 (clone HI100), CD38 BUV661 (clone HIT2), CD25 BUV496 (clone 2A3), ICOS BUV615 (clone DX29), CD71 BV605 (clone M-A712), CD3 BUV805 (clone UCHT1), BLIMP1 BV421 (clone 6D3), BCL6 RB705 (clone K112-91), and KI67 BUV395 (clone B56) were obtained from BD. FoxP3 AF700 (clone PCH101) was obtained from eBioscience. HIV-1 p24 RD1 (clone KC57) and HIV-1 p24 FITC (clone KC57) were obtained from Beckman Coulter.

### DNAscope ISH staining and Quantification Methods

We used immunofluorescence and *in situ* hybridization staining to assess the quantity and location of vDNA+ cells. We sectioned FFPE blocks at 5 µm onto Colorfrost Plus Microscope Slides (Fisher #1255017). We deparaffinized pairs of slides by baking at 60 °C for 1 hr, then immersing in two changes of xylene for 5 minutes each and then in two changes of 100% ethanol for 5 min each. After rinsing with nuclease-free water, heat-induced epitope retrieval (HIER) was performed by boiling slides in a Biocare Decloaker Chamber for 15 min at 110 °C.

Slides bound for brightfield microscopy had HIER in AR9 buffer (Akoya #AR900250ML diluted to 1x concentration in nuclease-free water), were immediately rinsed in room temperature nuclease-free water, and stained on a Leica BOND RX using an HIV clade B sense probe (ACD #425538) and RNAscope 2.5 LS Reagent Kit – Red (ACD #322150). They were then counterstained in CAT Hematoxylin (Biocare #CATHE-GL) for 1 min and blued in saturated sodium ammonium hydroxide for 1 min. Slides were covered in Clear-Mount (EMS #17985-15, diluted 50% v/v with water) and let dry at room temperature, then briefly immersed in xylene and cover slipped with Permount (ThermoFisher #SP15-500) and EMS Cover Glass – Gold Seal (thickness #1, #63768-10). These slides were the source of vDNA data.

Slides bound for fluorescent microscopy had HIER in in RNAscope Target Retrieval (ACD #322001 diluted to 1x concentration in nuclease-free water) and cooled in the retrieval solution for 20 min, then were rinsed in nuclease-free water and photobleached under a Sunny Light LED Therapy Lamp (OneSunrise, cat #SL-1A, color temperature 6500K) at maximum, notionally 32,000 lux, for 45 min. In this time, slides were covered in photobleach solution (12.84 mL PBS, 2.25 mL 30% H2O2, 69 µL 5 N NaOH). Immunofluorescent staining was performed by incubating slides for 1 hr at room temperature in a cocktail of antibodies, each at 1:100 concentration in 0.25% casein solution, against CD20 (clone L26, Biocare #CM004C) and CD4 (clone EPR6855, AbCam #ab133616). Signal was developed by incubating slides for 2 hr at room temperature in a cocktail of Donkey anti-Rabbit IgG (H+L) Highly Cross-Adsorbed Secondary Antibody, Alexa Fluor™ 488, and Donkey anti-Mouse IgG (H+L) Cross-Adsorbed Secondary Antibody, DyLight™ 755, (ThermoFisher #A21206 and #SA5-10171), each at 1:100 concentration in 0.25% casein solution. Nuclei were stained with DAPI (Invitrogen cat #D1306) at 1:10,000 in water for 10 minutes, cover slipping with Prolong Diamond Antifade Mountant (Invitrogen cat #P36970) and EMS Cover Glass – Gold Seal (thickness #1.5, #63791-01).

All slides were scanned on a Zeiss Axio Scan.Z1. Image analysis was performed using HALO/HALO-AI (Indica Labs) first on DNAscope stained slides we used HALO AI v3.6.4134, a Nuclei segmentation classifier that was trained to detect hematoxylin-stained nuclei. This was followed by using an additional classifer in HALO AI v4.1.5944.161 in an ISH v4.2.3 analysis module, detecting vDNA signal in whole lymph node annotations. Finally, we manually curated to exclude artifact chromagen signal and compared to the paired CD4-and-CD20-stained slide to determine whether vDNA+ cells were in a BCF, and can be inferred to be a T_FH_ cell, or located in the extrafollicular paracortical region. Figures were made in GraphPad Prism 10.5.0.

### Graphics

All figures were made in R with the following packages: ggplot2, Seurat, ArchR, ComplexHeatmap, and patchwork or are from FlowJo (v10, BD) unless otherwise stated. All code to produce graphs can be found in the study GitHub repository.

---

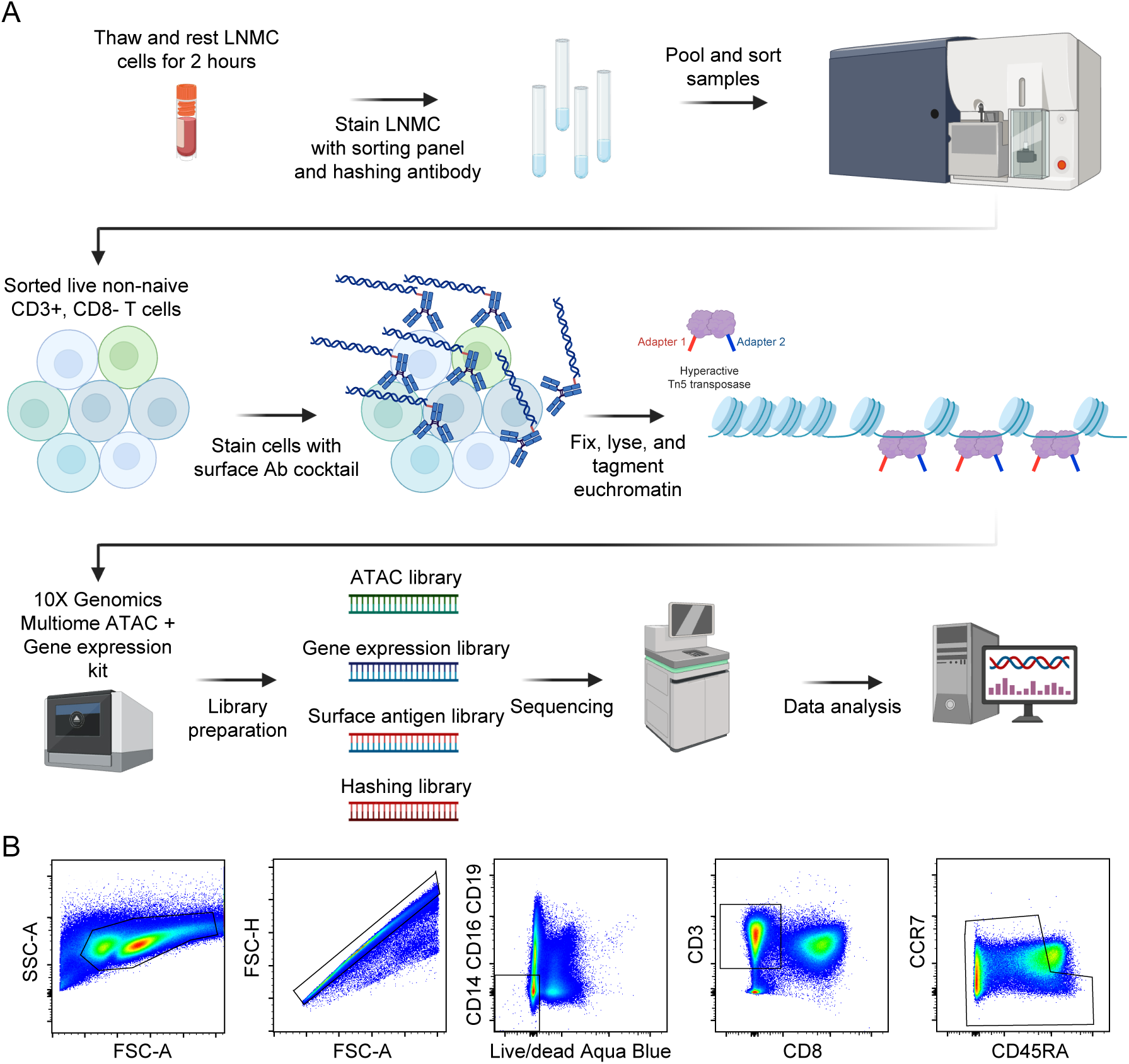

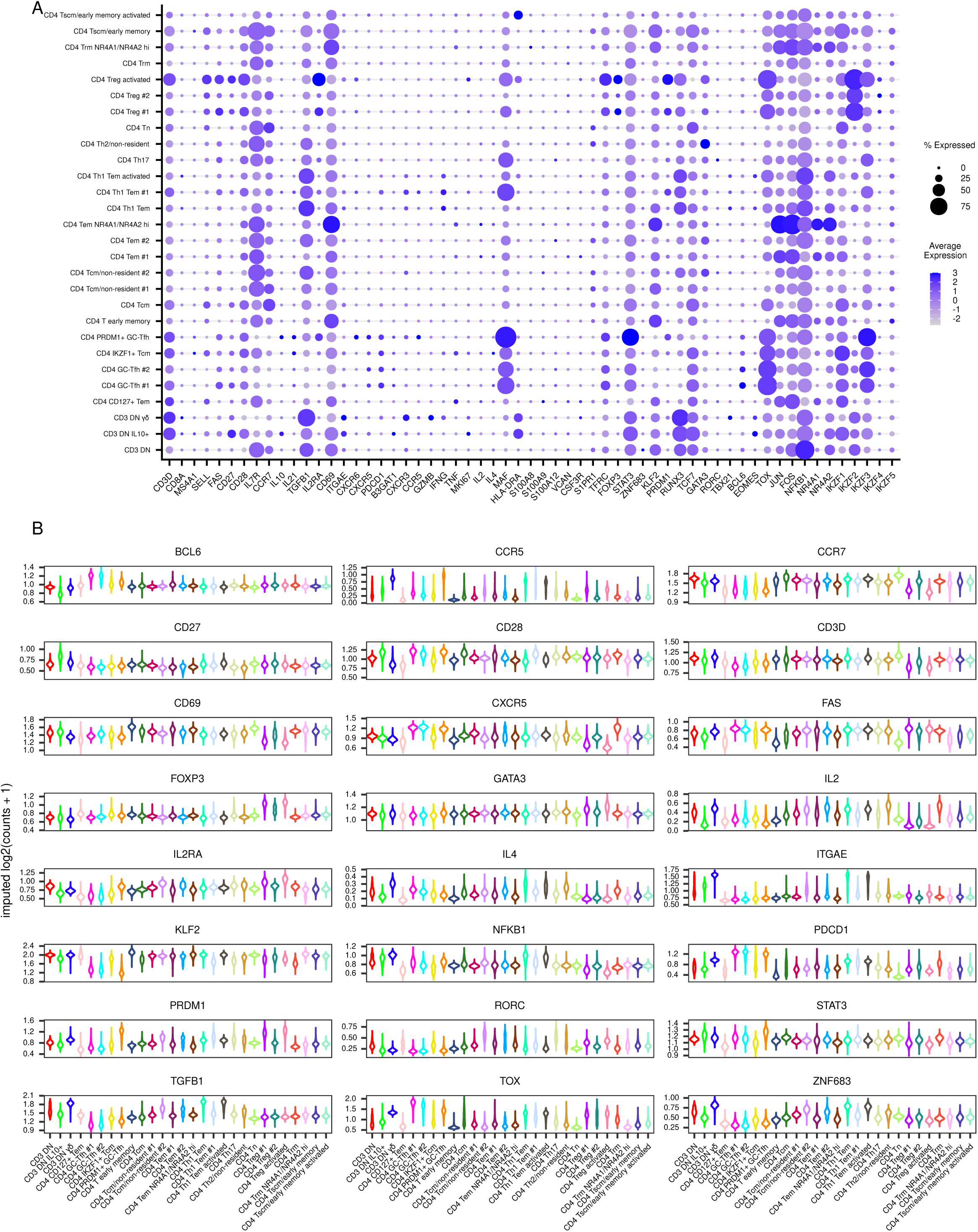

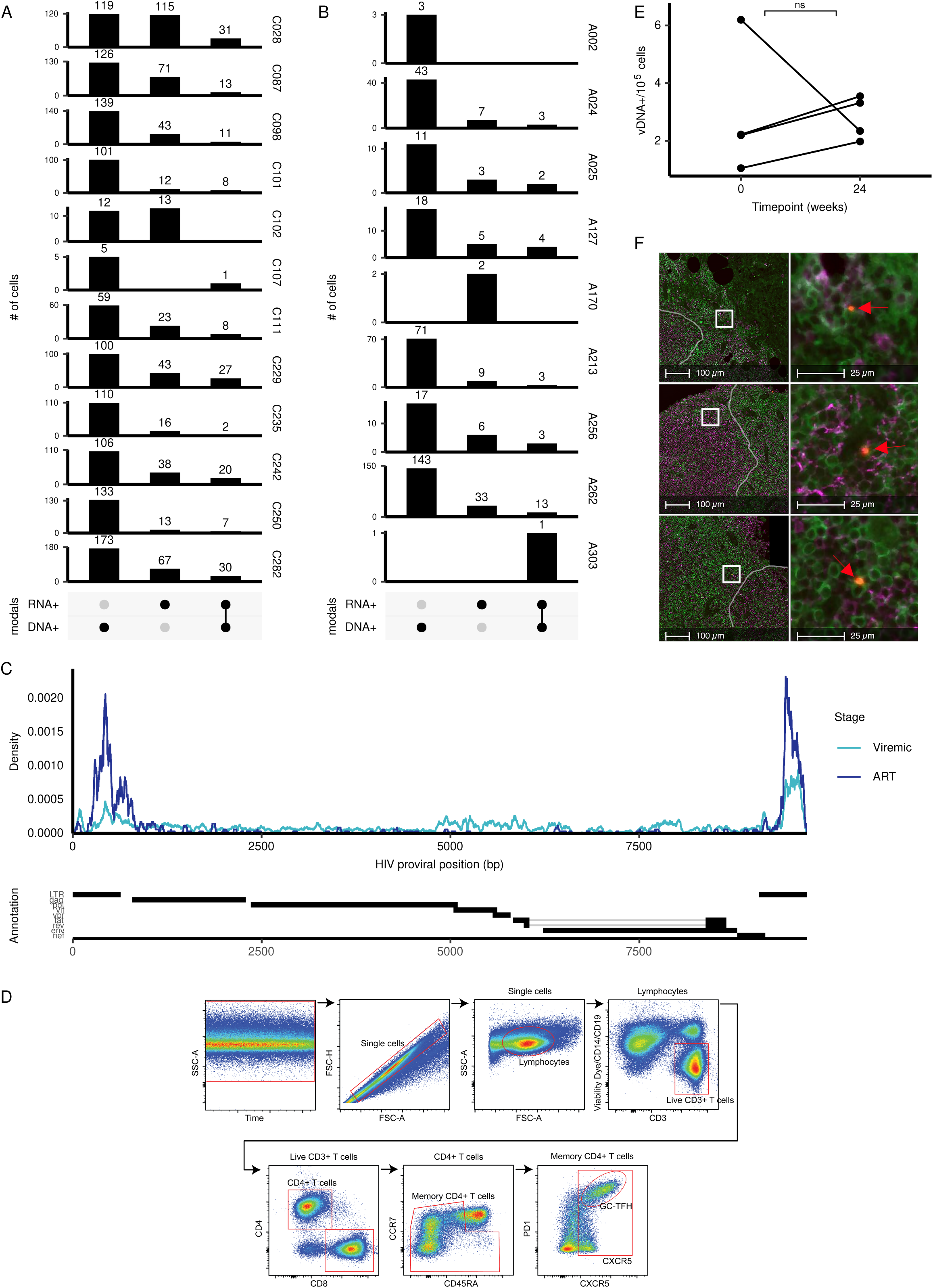

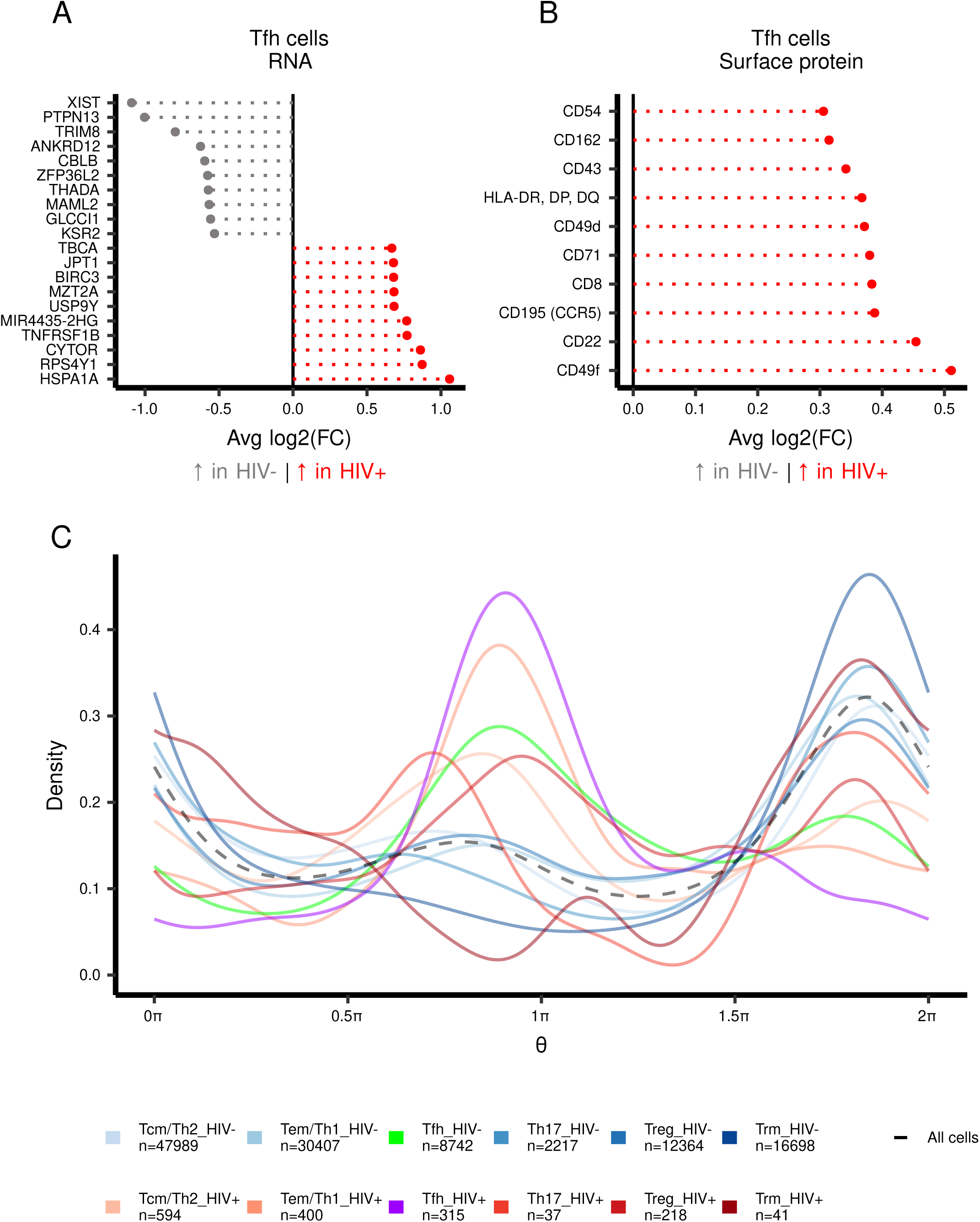

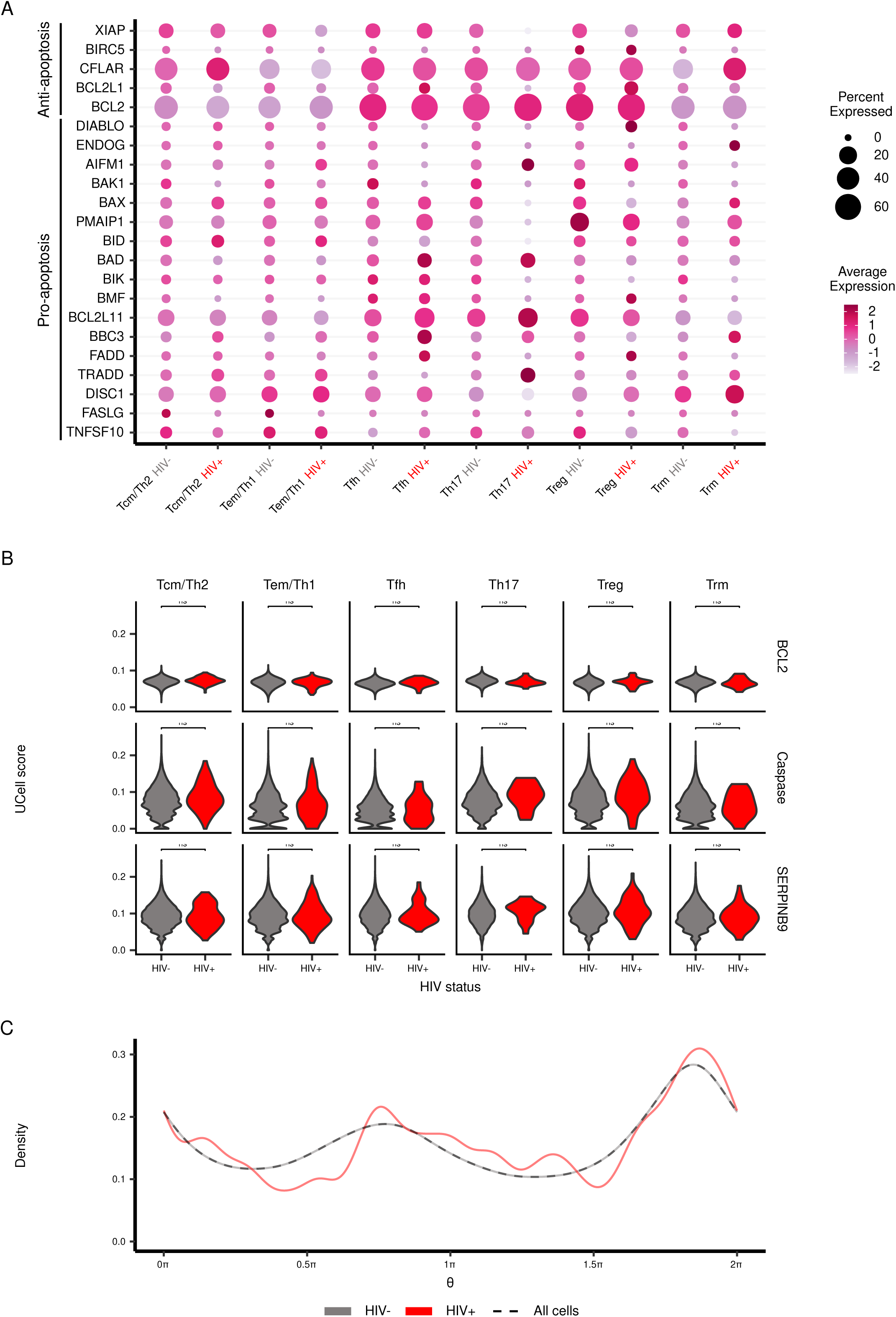

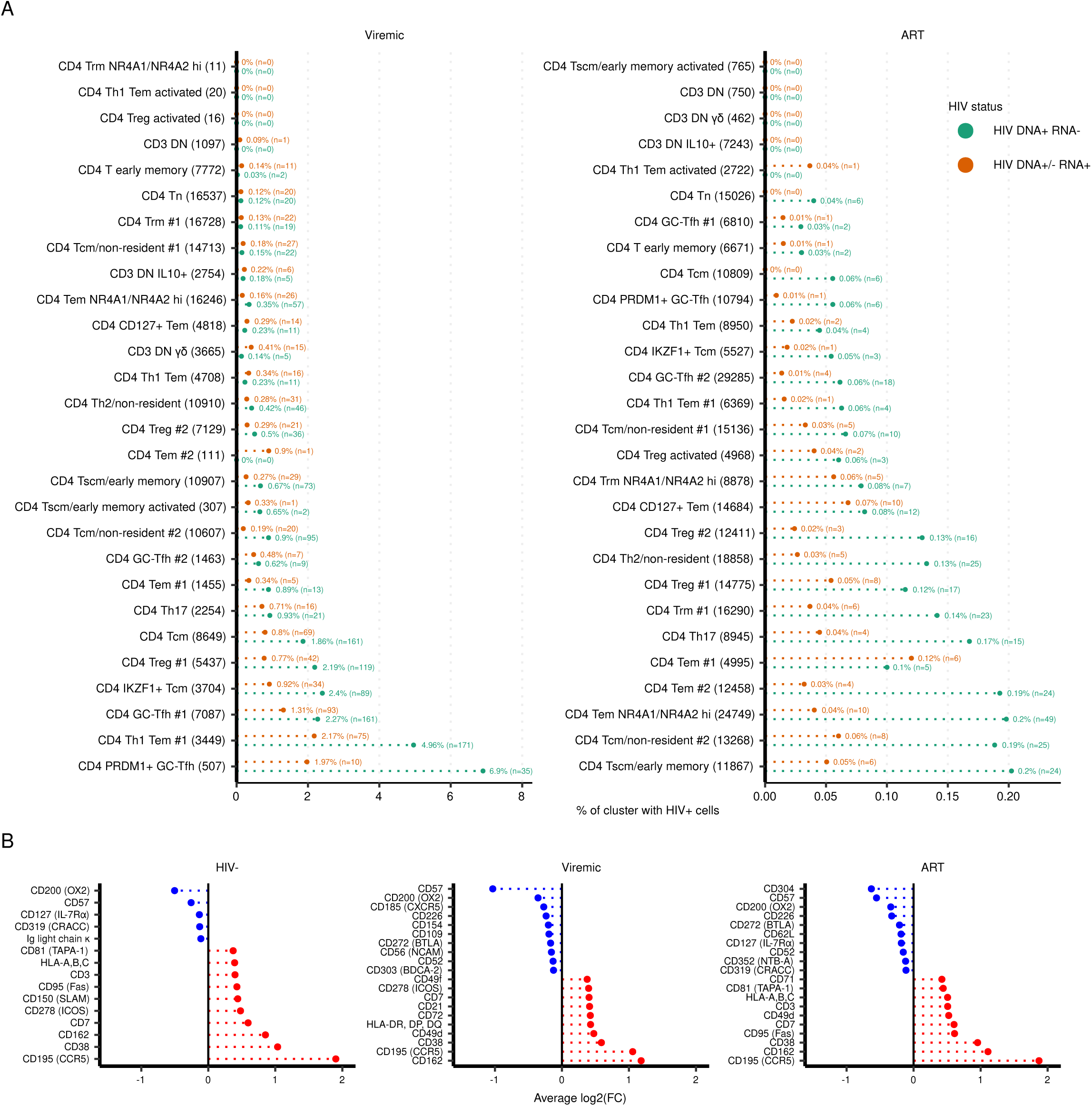

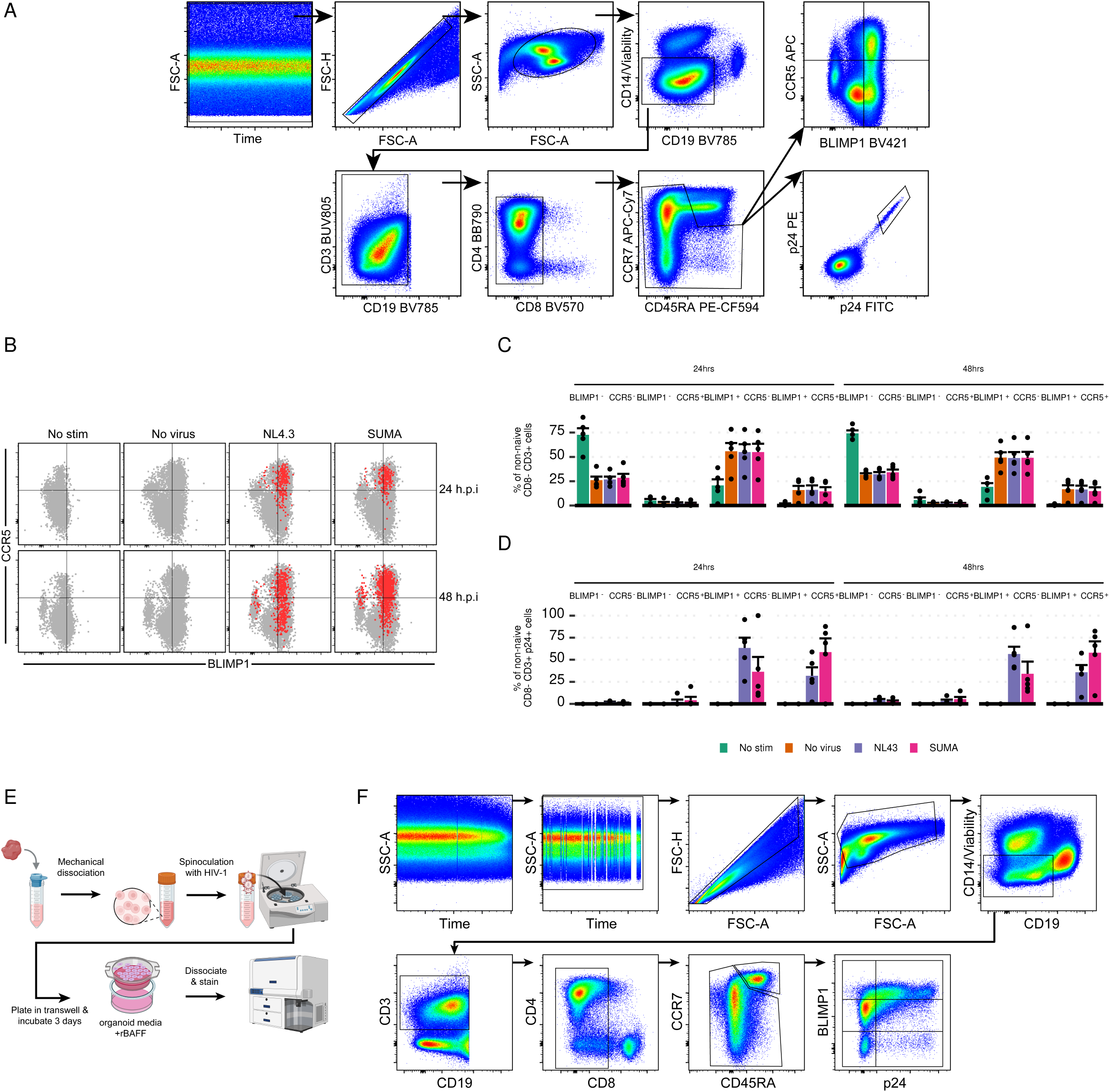

