## Supplemental Figures for "Selective infection and loss of PRDM1+ LN Tfh cells in uncontrolled HIV infection precludes formation of Tfh reservoirs under ART"

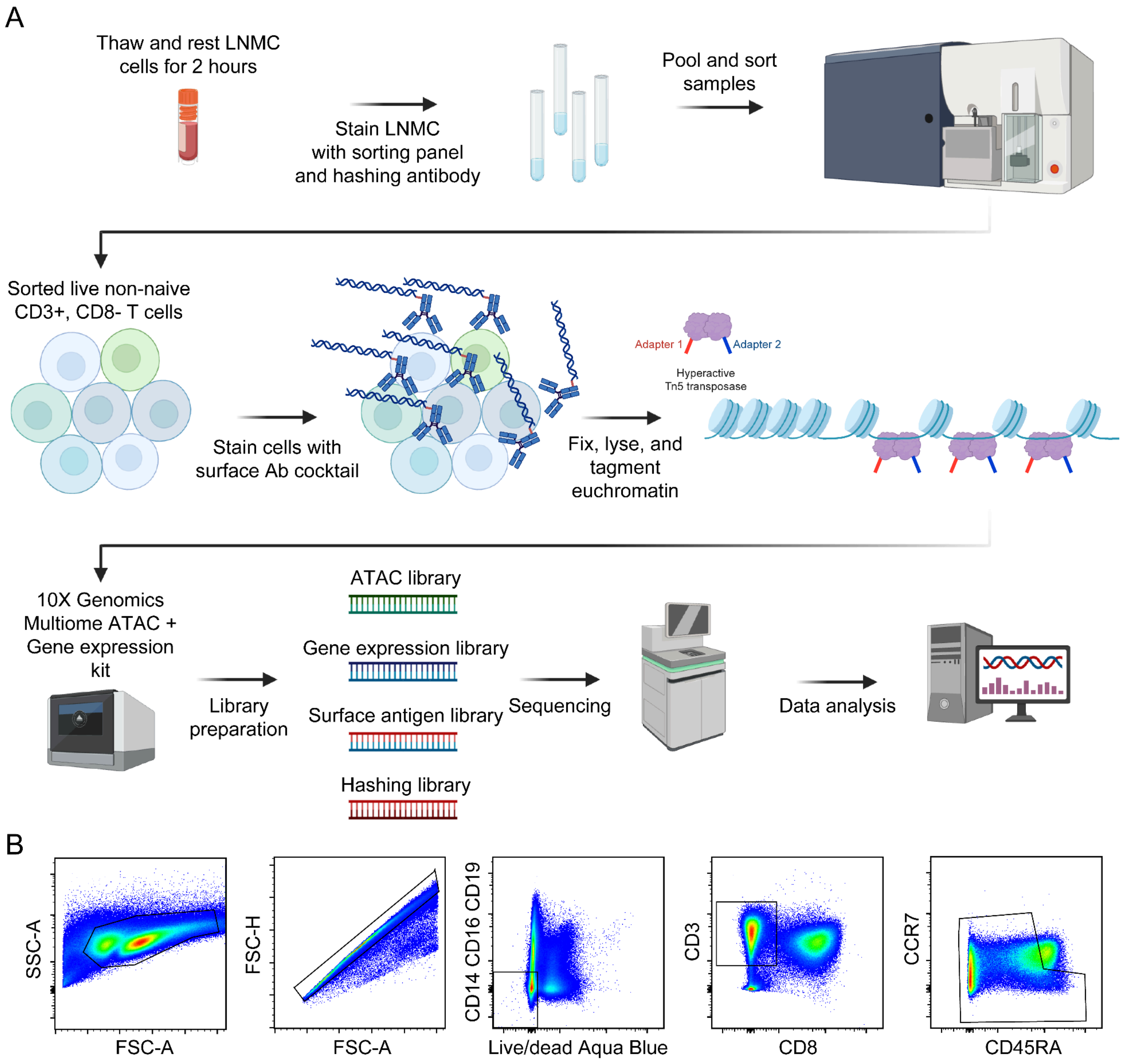


**Supplemental Figure 1:** (A) Schematic of the overall workflow for sample preparation starting from lymph node mononuclear cells (LNMC) to scDOGMAseq library preparation and sequencing. Graphics were made using biorender.com. (B) Representative sort strategy for the enrichment of non-naive CD3+ CD8- T cells. The gating hierarchy is shown from left to right.

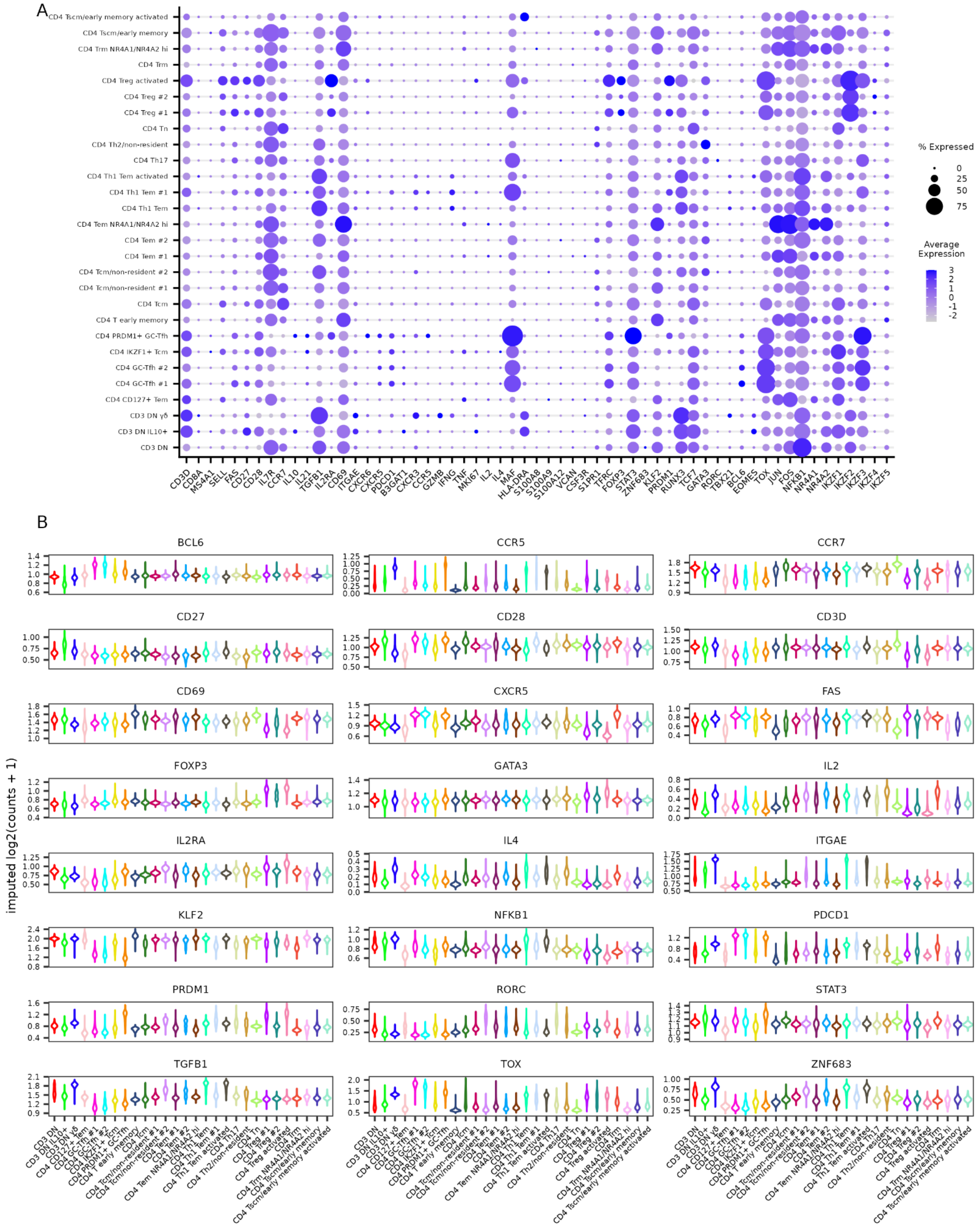
A
CD4 Tscm/early memory activated
CD4 Tscm/early memory
CD4 Trm NR4A1/NR4A2 hi
CD4 Trm
CD4 Treg activated
CD4 Treg #2
CD4 Treg #1
CD4 Tn
CD4 Th2/non-resident
CD4 Th17
CD4 Th1 Tem activated
CD4 Th1 Tem #1
CD4 Th1 Tem
CD4 Tem NR4A1/NR4A2 hi
CD4 Tem #2
CD4 Tem #1
CD4 Tcm/non-resident #2
CD4 Tcm/non-resident #1
CD4 Tcm
CD4 T early memory
CD4 PRDM1+ GC-Tfh
CD4 IKZF1+ Tcm
CD4 GC-Tfh #2
CD4 GC-Tfh #1
CD4 CD127+ Tem
CD3 DN γδ
CD3 DN IL10+
CD3 DN
CD3D CD8A MS4A1 SELL FAS CD27 CD28 IL7R CCR7 IL10 IL21 TGFB1 IL2RA CD69 ITGAE CXCR6 CXCR5 PDCD1 B3GAT1 CXCR3 CCR5 GZMB IFNG TNF MKI67 IL2 IL4 MAF HLA-DRA S100A8 S100A9 S100A12 VCAN CSF3R S1PR1 TFRC FOXP3 STAT3 ZNF683 KLF2 PRDM1 RUNX3 TCF7 GATA3 RORC TBX21 BCL6 EOMES TOX JUN FOS NFKB1 NR4A1 NR4A2 IKZF1 IKZF2 IKZF3 IKZF4 IKZF5
% Expressed
0
25
50
75
Average Expression
3
2
1
0
-1
-2
B
BCL6
CCR5
CCR7
CD27
CD28
CD3D
CD69
CXCR5
FAS
FOXP3
GATA3
IL2
IL2RA
IL4
ITGAE
KLF2
NFKB1
PDCD1
PRDM1
RORC
STAT3
TGFB1
TOX
ZNF683
imputed log2(counts + 1)
CD3 DN
CD3 DN IL10+
CD3 DN γδ
CD4 CD127+ Tem
CD4 GC-Tfh #1
CD4 GC-Tfh #2
CD4 IKZF1+ Tcm
CD4 PRDM1+ GC-Tfh
CD4 T early memory
CD4 Tcm
CD4 Tcm/non-resident #1
CD4 Tcm/non-resident #2
CD4 Tem #1
CD4 Tem #2
CD4 Tem NR4A1/NR4A2 hi
CD4 Th1 Tem
CD4 Th1 Tem #1
CD4 Th1 Tem activated
CD4 Th17
CD4 Th2/non-resident
CD4 Tn
CD4 Treg #1
CD4 Treg #2
CD4 Treg activated
CD4 Trm
CD4 Trm NR4A1/NR4A2 hi
CD4 Tscm/early memory
CD4 Tscm/early memory activated

**Supplemental Figure 2:** Main panel of markers used for annotation from different modalities. (A) Average RNA expression (scaled normalized counts). (B) Imputed gene activity score from the ATAC modality (calculated by ArchR and imputed using MAGIC). Values represent log2(normalized counts + 1).

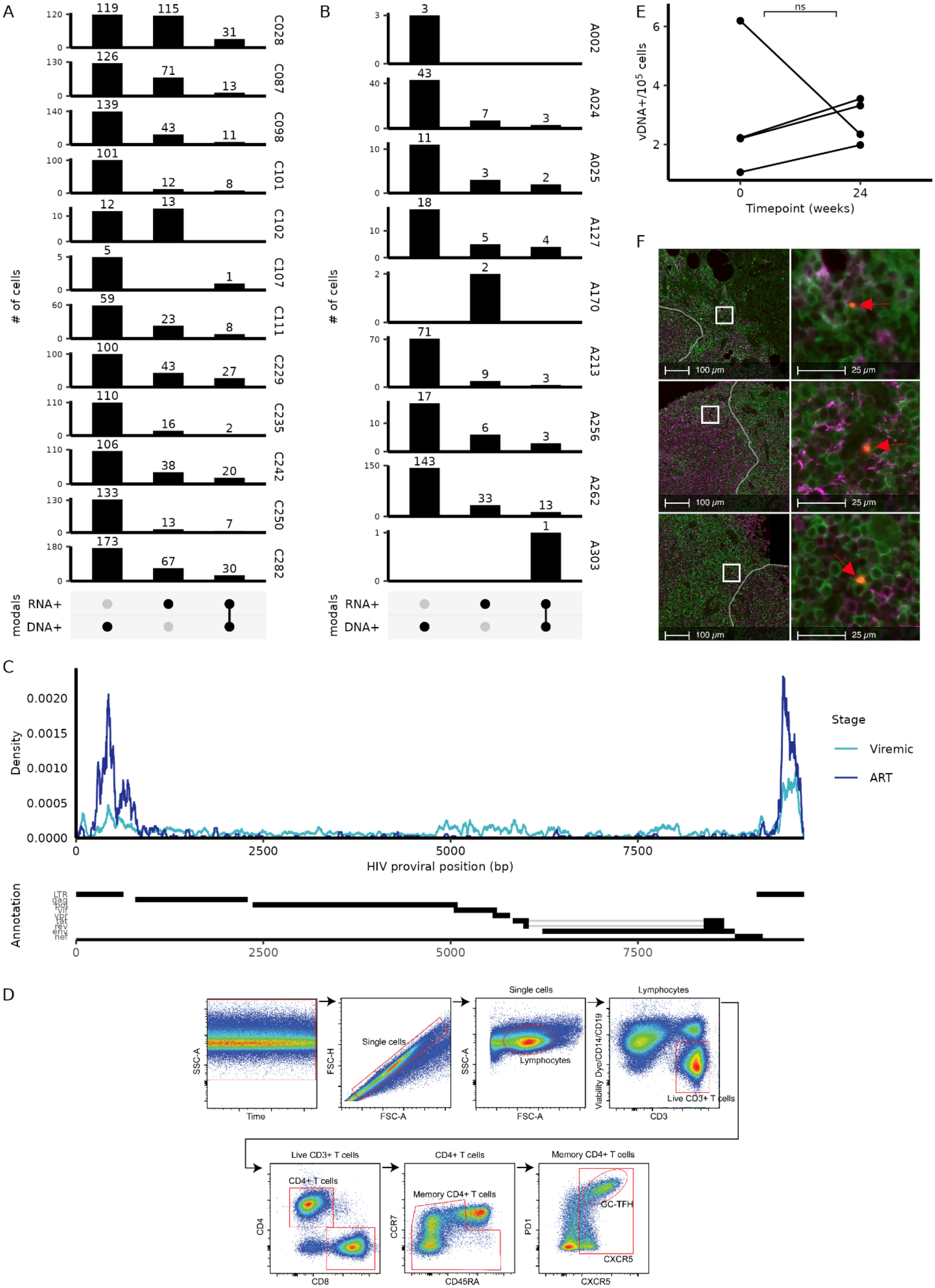

A
B
C
D
E
F
### of cells
modals
RNA+
DNA+
C028
C087
C098
C101
C102
C107
C111
C229
C235
C242
C250
C282
A002
A024
A025
A127
A170
A213
A256
A262
A303
ns
vDNA+/10^5 cells
Timepoint (weeks)
100 μm
25 μm
Density
HIV proviral position (bp)
Stage
Viremic
ART
Annotation
LTR
gag
pol
vif
vpr
tat
rev
env
nef
Single cells
Lymphocytes
Live CD3+ T cells
CD4+ T cells
Memory CD4+ T cells
GC-TFH
SSC-A
FSC-H
FSC-A
Time
Viability Dye/CD14/CD19
CD3
CD4
CD8
CCR7
CD45RA
PD1
CXCR5

**Supplemental Figure 3:** (A) Counts of cells by viremic individuals with detection of HIV DNA and/or HIV RNA. (B) Counts of cells by ART-treated individuals with detection of HIV DNA and/or HIV RNA. (C) Density coverage of reads from the ATAC modality across the HIV genome. (D) Representative gating strategy for CD4+ GC-Tfh cells by flow cytometry. (E) Percentage of viral DNA+ cells detected in each tissue sample by HIV DNAscope. Wilcoxon Rank Sum Test was performed with Bonferroni multiple test adjustment. ns = not significant (adjusted p value >= 0.05). (F) Representative immunofluorescence microscopy image on lymph nodes. Each row represents a separate area of the tissue slice. Left column represents a lower magnification capture. White squares denote an inset that is then magnified and shown in the right column. Green = CD4, Red = vDNA, Magenta = CD20, Grey lines = BCF boundary.

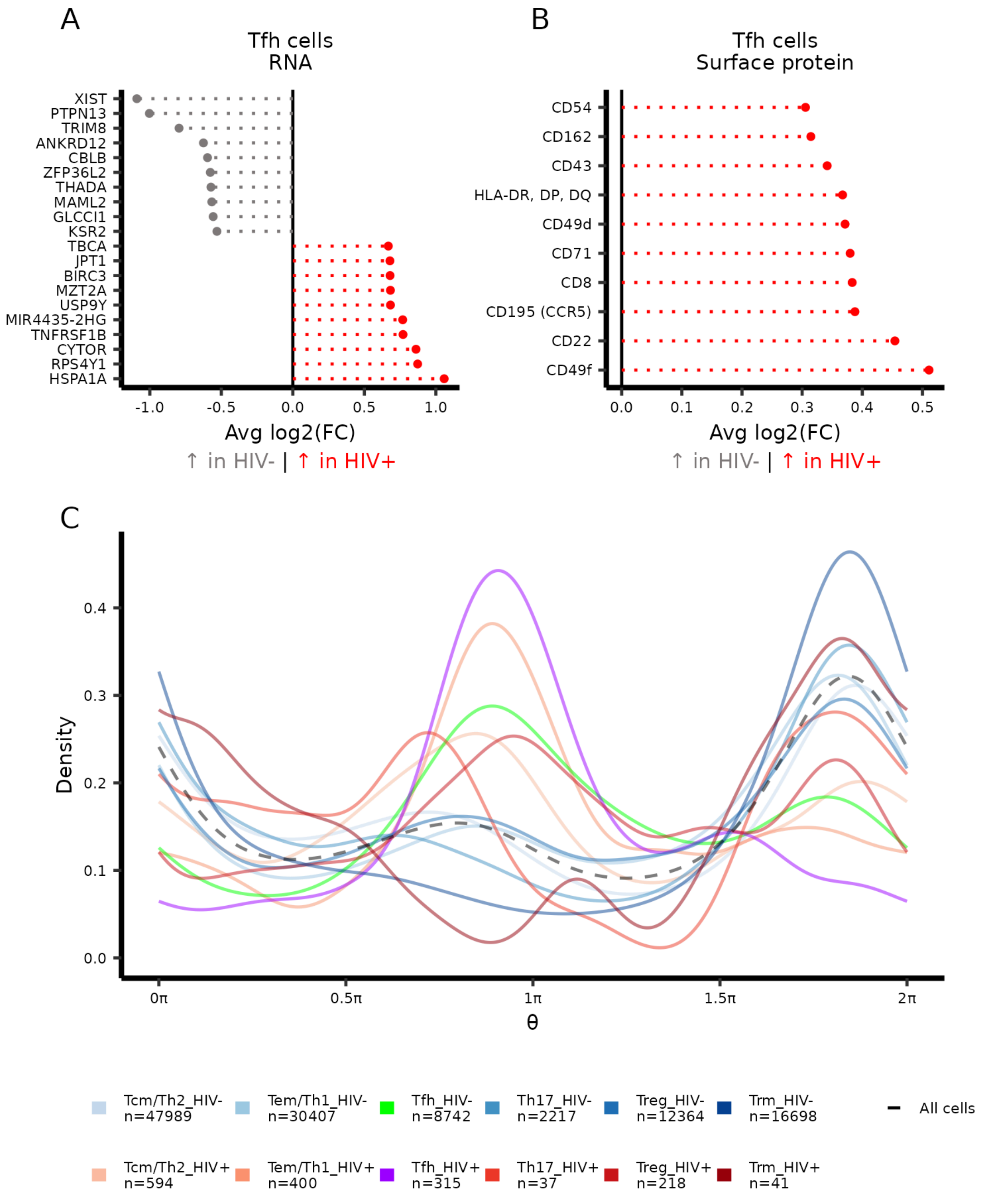


Supplemental Figure 4: (A and B) Differential RNA and surface protein expression between HIV+ and HIV- cells. Markers that are positively enriched in HIV+ cells are shown in red (positive fold change) while markers that are enriched in the HIV- subset are shown in blue (negative fold change). Only the top 10 (by fold change) significant (Wilcoxon Rank Sum followed by Bonferroni-Hochberg correction; adjusted p value < 0.05) markers are shown in each direction. Markers with at least 30% expression in one comparison group were tested. (C) Cell cycle position ($\theta$ where $\pi$ is approximately G2M and $1.5\pi$ is middle of M stage) was estimated with the tricycle package to compare between HIV+ cells and HIV- cells when separated by cell subset. The black dashed line represents the distribution of all cells from viremic PWH.

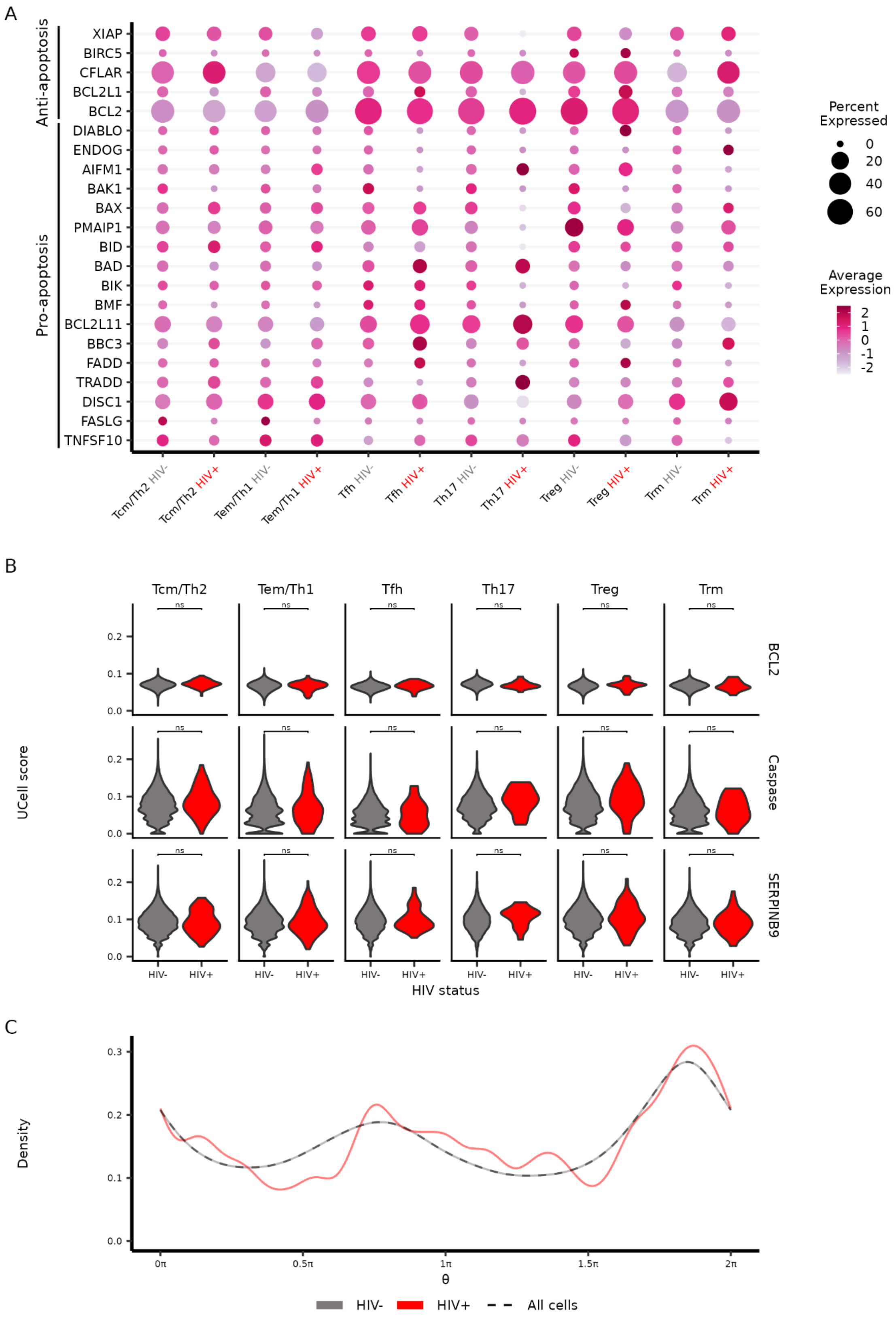


**Supplemental Figure 5:** (A) Dot plot showing scaled RNA expression levels of various pro- and anti-apoptotic genes (O'Brien and Kirby 2008) in ART-treated PWH separated by HIV+ and HIV- cells within cell subsets. (B) UCell scores were calculated for various gene sets from Harmonizome 3.0 (Diamant et al. 2025): BCL2 (“bcl2” from the dataset “GeneRIF Biological Term Annotations”); caspase (“caspase cascade in apoptosis” from the dataset “Biocarta Pathways”); and SERPINB9 (“SERPINB9” from the dataset “Pathway Commons

Protein-Protein Interactions") (C) Cell cycle position (θ where π is approximately G2M and 1.5π is middle of M stage) was estimated with the tricycle package to compare between aggregated HIV+ cells and HIV- cells. The black dashed line represents the distribution of all cells from ART-treated PWH.

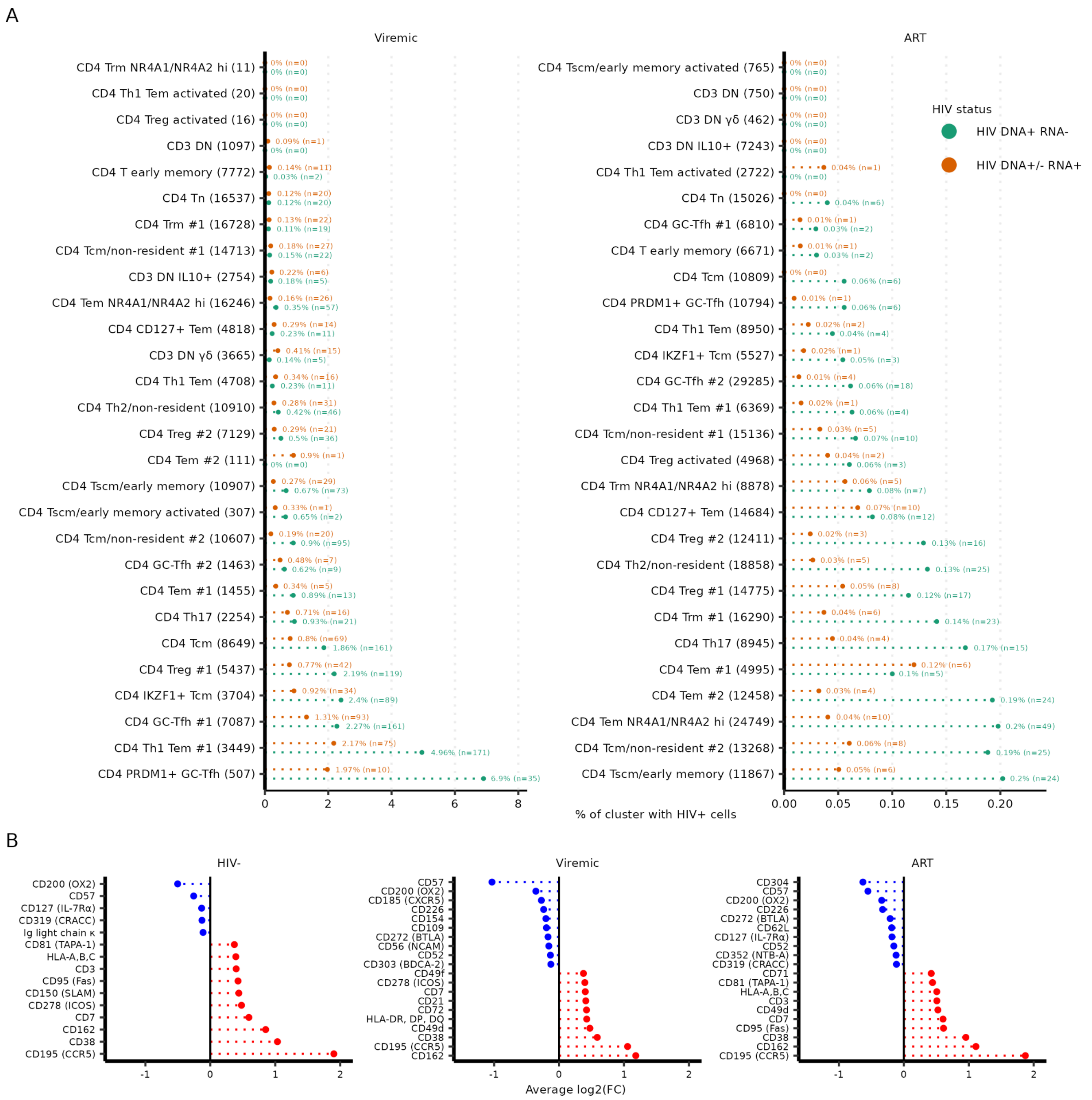


**Supplemental Figure 6:** (A) Aggregate percentage of cells within a cluster (separated by HIV infection stage) that has either HIV DNA+ RNA- (green) or HIV DNA+/- RNA+ (orange) cells. Clusters are arranged separately for viremic and ART in ascending order by percentage of HIV+ cells (both HIV DNA+ RNA- and HIV DNA+/- RNA+) in a cluster. (B) Differential surface protein expression between the CD4 PRDM1+ GC-Tfh subset and PRDM1- GC-Tfh subsets (CD4 GC-Tfh #1 and CD4 GC-Tfh #2). Markers that are positively enriched in the PRDM1+ subset are shown in red (positive fold change) while markers that are enriched in the PRDM1- subset are shown in blue (negative fold change). Only the top 10 (by fold change) significant (Wilcoxon Rank Sum followed by Bonferroni-Hochberg correction; adjusted p value < 0.05) markers are shown in each direction. Markers with at least 30% expression in one comparison group were tested.

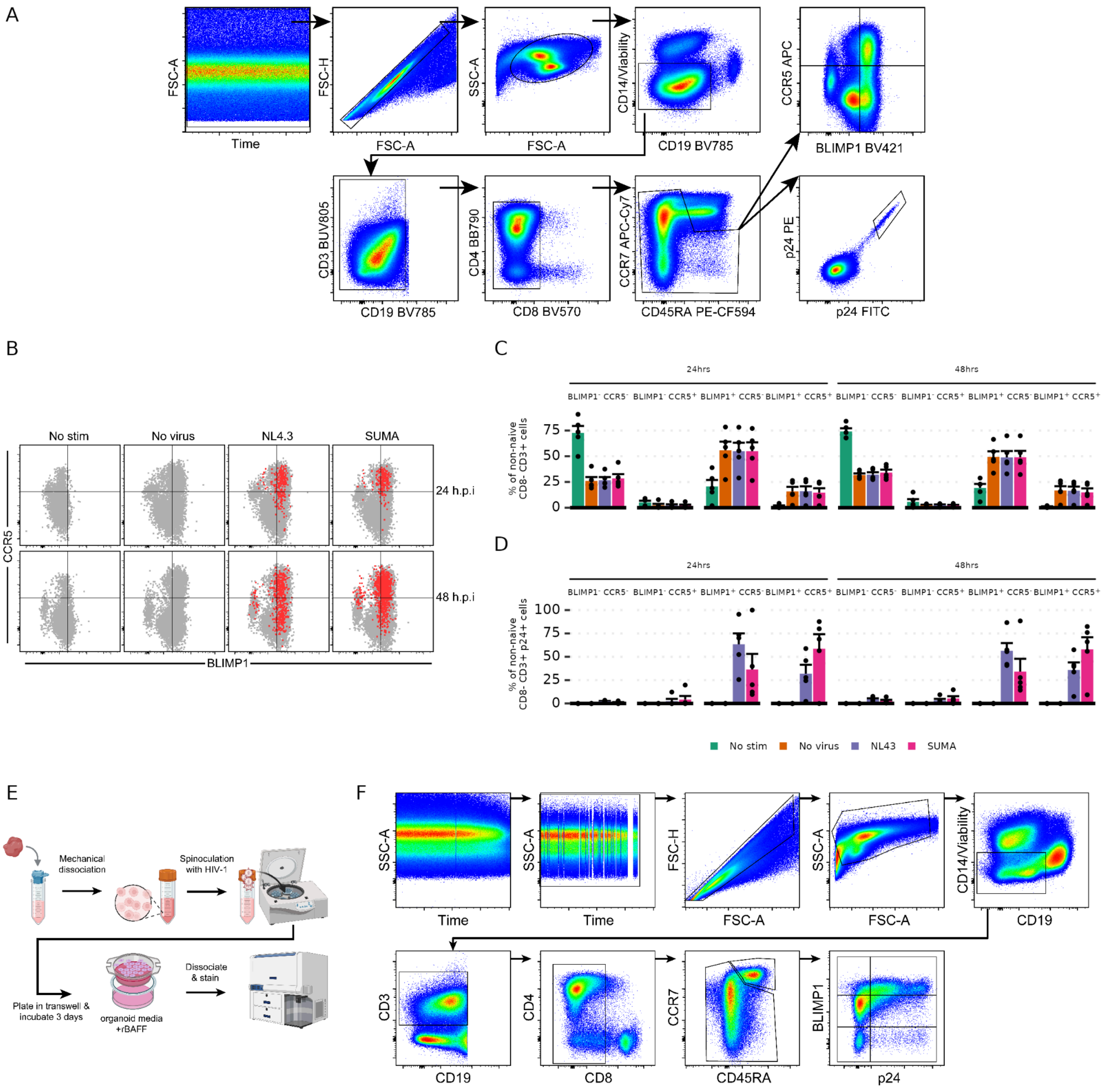


**Supplemental Figure 7:** (A) Gating strategy for in vitro infection of activated CD4 T cells. A representative infection with SUMA (MOI = 0.1) at 48 hours post infection (h.p.i) is shown. (B) Non-naive CD8- T cells are shown in grey while p24+ (double-positive with two separate fluorochromes as shown in (A)) are shown in red. No virus, NL4.3, and SUMA conditions have undergone stimulation. (C) Proportions of non-naive CD8- T cells that are found in the combinatorial BLIMP1 and CCR5 gates. Bars are colored based on virus tropism and negative controls. (D) Same as (C) but showing non-naive CD8- p24+ cells. (E) Schematic for tonsil organoid experiment. Diagrams were made with biorender.com (F) Representative gating strategy for the tonsil organoid experiments to gate for non-naive CD8- T cells with combinatorial gates for BLIMP1 and p24 expression.
