## Supplementary material for "Selective infection and loss of PRDM1+ LN Tfh cells in uncontrolled HIV infection precludes formation of Tfh reservoirs under ART": Main Table 1

Table 1: Donor information

| Stage | Donor | Sex | Age (years) | Tissue | CD4 count (cells/ul) | Duration of ART (months) | Viral Load (copies/ml) |
|---|---|---|---|---|---|---|---|
| ART | A002 | Male | 32 | CLN | 956 | 16 | Below LOD (<40) |
| | A024 | Male | 37 | ILN | 583 | 26 | Below LOD (<40) |
| | A025 | Male | 28 | ILN | 601 | 16 | Below LOD (<40) |
| | A127 | Male | 32 | CLN | 1104 | 25 | Below LOD (<40) |
| | A170 | Male | 29 | CLN | 390 | 25 | Below LOD (<40) |
| | A213 | Male | 40 | CLN | 753 | 241 | Below LOD (<40) |
| | A223 | Male | 42 | CLN | 674 | 57 | Below LOD (<40) |
| | A256 | Male | 40 | CLN | 322 | 32 | Below LOD (<40) |
| | A262 | Male | 33 | CLN | 437 | 36 | Below LOD (<40) |
| | A266 | Male | 29 | CLN | 849 | 56 | Below LOD (<40) |
| | A293 | Female | 28 | CLN | 672 | 69 | Below LOD (<40) |
| | A303 | Male | 20 | CLN | 507 | 11 | Below LOD (<40) |
| | A304 | Male | 26 | CLN | 649 | 22 | Below LOD (<40) |
| Viremic | C021 | Male | 28 | CLN | 386 | - | 10,620 |
| | C028 | Female | 38 | CLN | 531 | - | 25,951 |
| | C052 | Male | 33 | CLN | 499 | - | 1,158,541 |
| | C087 | Male | 32 | CLN | 462 | - | 1,771,593 |
| | C094 | Male | 23 | CLN | 509 | - | 143,916 |
| | C098 | Male | 21 | CLN | 251 | - | 1,019,989 |
| | C101 | Male | 31 | ILN | 281 | - | 1,935,095 |
| | C102 | Male | 26 | CLN | 410 | - | 1,692,571 |
| | C107 | Male | 33 | CLN | 368 | - | 6,326,452 |
| | C111 | Male | 22 | CLN | 507 | - | 552,134 |
| | C121 | Male | 23 | CLN | 341 | - | 343,598 |
| | C229 | Male | 18 | CLN | 265 | - | 3,882,624 |
| | C235 | Male | 37 | ILN | 320 | - | 1,067,647 |
| | C242 | Male | 20 | CLN | 174 | - | 715,509 |
| | C250 | Male | 29 | ILN | 545 | - | 259,708 |
| | C282 | Male | 25 | CLN | 812 | - | 3,087,470 |
| HIV- | U190 | Male | 30 | CLN | 1148 | - | Below LOD (<40) |
| | U191 | Male | 37 | CLN | 1521 | - | Below LOD (<40) |
| | U287 | Male | 29 | ILN | 1666 | - | Below LOD (<40) |
| | U294 | Male | 30 | ILN | 420 | - | Below LOD (<40) |
