## Supplemental Table 1 for "Selective infection and loss of PRDM1+ LN Tfh cells in uncontrolled HIV infection precludes formation of Tfh reservoirs under ART"

**Supplemental Table 1:** Curated list of genes for annotation.

| Gene | Notes |
|---|---|
| CD3D | Lineage - T cell |
| CD8A | Lineage - CD8 T cell |
| MS4A1 | Lineage - B cell |
| SELL | T - Memory |
| FAS | T - Memory |
| CD27 | T - Memory |
| CD28 | T - Memory |
| IL7R | T - Memory |
| CCR7 | T - Memory |
| IL10 | Treg |
| IL21 | Tfh functionality |
| TGFB1 | Treg |
| IL2RA | Treg/activation |
| CD69 | T - Activation/residency |
| ITGAE | T - Residency |
| CXCR6 | T - Residency |
| CXCR5 | Follicular marker/trafficking |
| PDCD1 | T - Inhibitory |
| B3GAT1 | T/NK – Effector/differentiation |
| CXCR3 | Activation/trafficking |
| CCR5 | Activation |
| GZMB | T - Effector |
| IFNG | T - Effector |
| TNF | T - Effector |
| MKI67 | Proliferation/activation (gene for Ki67) |
| IL2 | T - Functionality |
| IL4 | Th2 |
| MAF | M2 macrophages/anti-inflammation |
| HLA-DRA | Multi purpose (activation + monocyte) |
| S100A8 | Monocyte from Mulder et al., Immunity 2021 |
| S100A9 | Monocyte from Mulder et al., Immunity 2021 |
| S100A12 | Monocyte from Mulder et al., Immunity 2021 |
| VCAN | Monocyte from Mulder et al., Immunity 2021 |
| CSF3R | Monocyte from Mulder et al., Immunity 2021 |
| S1PR1 | Residency |
| TFRC | Proliferation/activation (gene for CD71) |
| FOXP3 | Treg |
| STAT3 | Th17 |
| ZNF683 | T - Residency/effector |
| KLF2 | T - Effector/trafficking |
| PRDM1 | T - Nonclassical Tfh (gene for Blimp1) |
| RUNX3 | T - Residency/development |
| TCF7 | T - Memory |
| GATA3 | Th2 |
| RORC | Th17 |
| TBX21 | T - Memory |
| BCL6 | Tfh driver |
| EOMES | T - Memory |
| TOX | Inhibitory/exhaustion |
| JUN | T - Activation |
| FOS | T - Activation |
| NFKB1 | T - Activation |
| NR4A1 | T - Recent TCR stim |
| NR4A2 | T - Recent TCR stim |
| IKZF1 | IKZ family - 1 Ikaros |

| | |
|---|---|
| IKZF2 | IKZ family - 2 Helios |
| IKZF3 | IKZ family - 3 Aiolos |
| IKZF4 | IKZ family - 4 Eos |
| IKZF5 | IKZ family - 5 Pegasus |
| SELL | T - Memory |
| FAS | T - Memory |
| CD27 | T - Memory |
| CD28 | T - Memory |
